# Complex Conformational Space of RNA Polymerase II C-Terminal Domain upon Phosphorylation

**DOI:** 10.1101/2023.04.20.537737

**Authors:** Weththasinghage D. Amith, Bercem Dutagaci

## Abstract

Intrinsically disordered proteins (IDPs) have been closely studied during the past decade due to their importance for many biological processes. The disordered nature of this group of proteins makes it difficult to observe its full span of the conformational space either using experimental or computational studies. In this article, we explored the conformational space of the C-terminal domain (CTD) of RNA polymerase II (Pol II), which is also an intrinsically disordered low complexity domain, using enhanced sampling methods. We provided a detailed conformational analysis of model systems of CTD with different lengths; first with the last 44 residues of the human CTD sequence and finally the CTD model with two heptapeptide repeating units. We then investigated the effects of phosphorylation on CTD conformations by performing simulations at different phosphorylated states. We obtained broad conformational spaces in non-phosphorylated CTD models and phosphorylation has complex effects on the conformations of the CTD. These complex effects depend on the length of the CTD, spacing between the multiple phosphorylation sites, ion coordination and interactions with the nearby residues.

## INTRODUCTION

During the last decades, intrinsically disordered proteins (IDPs) have been recognized as an important class of proteins due to their relevance to many biological processes.^1, 2^ Cell signaling and regulation^3^, stress response^4^, human neurodegenerative diseases^5^ and cellular liquid-liquid phase separation (LLPS)^6, 7^ are some of the biological phenomena which are associated with IDPs. One of the most challenging aspects of IDPs is to determine the full span of its conformational space using currently available experimental techniques such as small-angle X-ray scattering (SAXS), nuclear magnetic resonance (NMR) and circular dichroism (CD) spectroscopy. Such experimental methods have provided information on structural and dynamics features of IDPs, such as backbone conformations, secondary structures, overall shape and size of the molecules.^8–10^ However, converting these features to actual conformational ensembles remains to be a challenge. Therefore, the common approach became to develop and apply computational methods to obtain conformational spaces of IDPs and to use experimental features for assisting and/or validating the computational models. These computational methods include Monte Carlo approaches to generate ensembles^11, 12^, approaches based on molecular dynamics (MD) simulations by applying fragmentation of long IDP sequences^13, 14^ or enhanced sampling simulations in atomic or coarse-grained details.^5, 15–18^ Generative machine learning models were also developed, which utilized conformations obtained from MD simulations for training.^19–21^ Among computational methods, MD simulations with enhanced sampling techniques appeared to be a powerful method especially for relatively short sequences to provide valuable insights in atomic level interactions of IDPs in order to obtain an accurate conformational space.^15, 18, 22–25^

The C-terminal domain (CTD) of RNA polymerase II (Pol II) is a low complexity domain that contains heptapeptide (YSPTSPS) repeating sequence with the number of repeating units differs according to the organism.^26^ Pol II CTD is also recognized as an IDP due to its lack of defined secondary structure.^26^ Also, recent studies have shown Pol II involvement in LLPS formation^27, 28^ and suggested that CTD may play a fundamental role in such phase separation events. The length and phosphorylation pattern of CTD are also shown to have effects on LLPS formation.^27, 28^ Therefore, determination of an accurate conformational space for CTD upon phosphorylation is significantly important to recognize the structural features which would impact the Pol II CTD related LLPS inside a cell. Our knowledge on conformational analysis of CTD and structural effects of phosphorylation is limited as there are only a few experimental^26, 29–34^ and computational^22, 35, 36^ studies on the structure of CTD. Hence, in this work, we studied the conformational landscape of model systems of CTD and effects of phosphorylation on the conformations. We analyzed the CTD models using an enhanced sampling method, replica-exchange molecular dynamics (REMD)^37^ simulations, in order to sample a wide range of probable conformations. We applied REMD simulations on two model systems; one was the 44-residue tail of human CTD, which has available experimental data to validate our simulations,^22, 29, 30^ and the other was a peptide with a sequence of 2-heptapeptide repeat of CTD (2CTD). Simulations were performed on non-phosphorylated and phosphorylated CTD sequences to observe the effects of phosphorylation pattern on the conformations of CTD models. Phosphorylation introduced conformational changes to both CTD models with 44 residues and 2-heptapeptide repeats compared to their non-phosphorylated states; however, the phosphorylated models of 2-heptapeptide repeats showed complex effects on their conformational space, while 44-residue models showed somewhat expected conformational changes depending on their net charge which we will elaborate in detail under the results and discussion sections below.

## COMPUTATIONAL METHODS

### System preparation and equilibration

All the initial structures of CTDs were generated by using the Multiscale Modeling Tools in Structural Biology (MMTSB) toolset^38^, CHARMM^39^ package and MODELLER program.^40^ Then the solvated systems were prepared by using the CHARMM-GUI server^41–43^ with most extended initial model structures of CTDs from the previous step. For all CTD sequences, N-terminus and C-terminus were capped with acetyl (ACE) and -NHCH3 (CT3) groups, respectively. Table 1. shows all the CTD sequences prepared for this study.

**Table 1.**
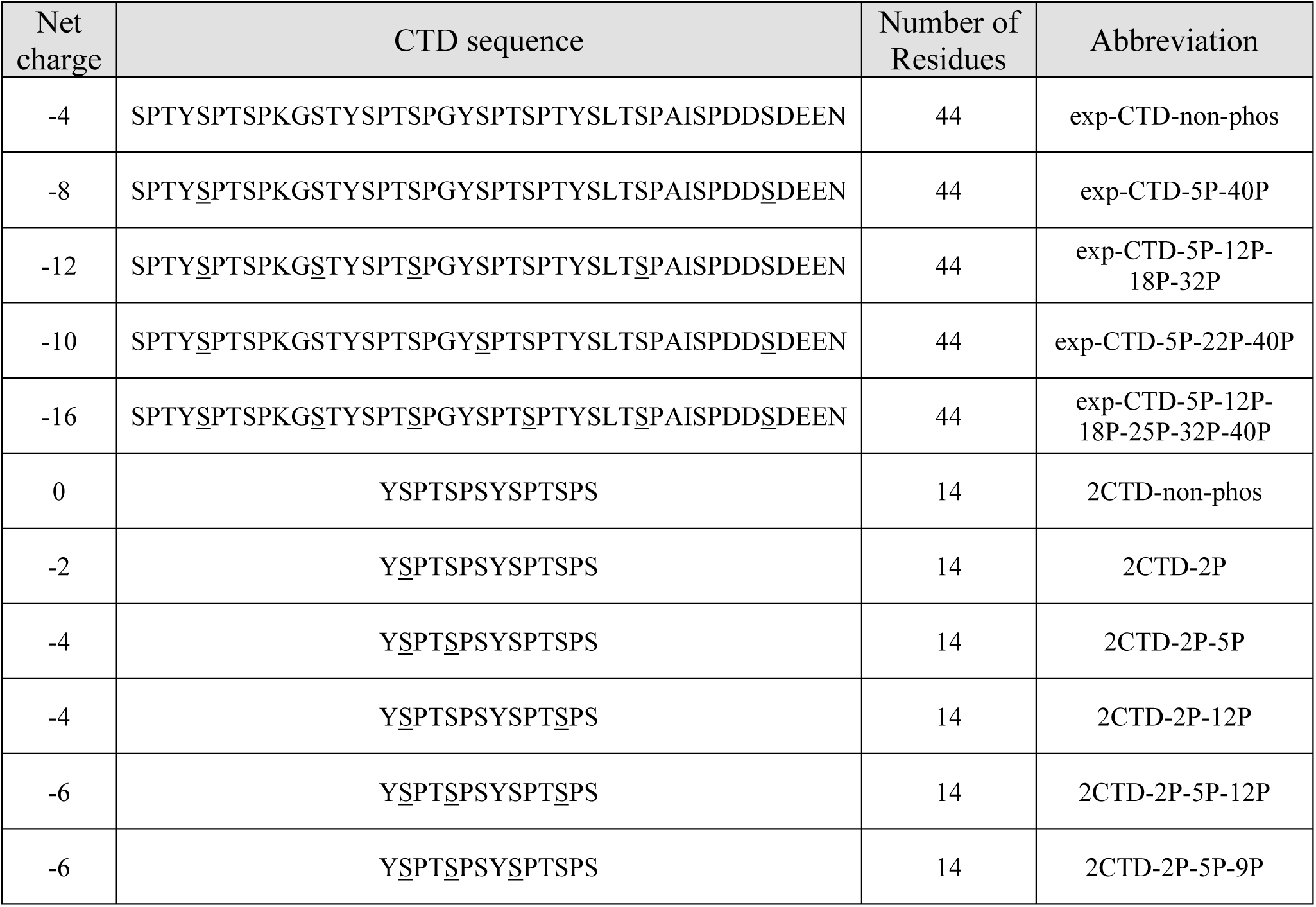
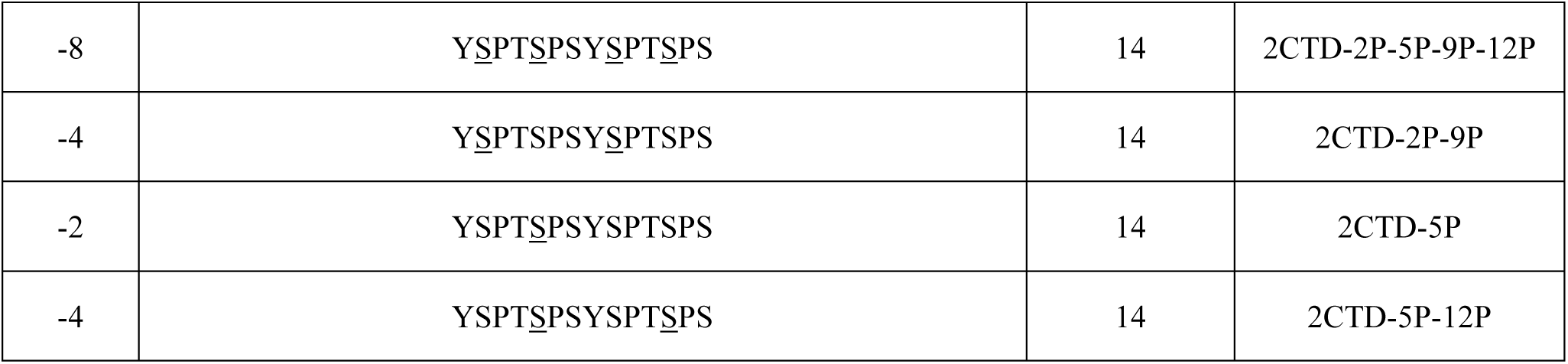
The CTD sequences, their net charges, number of residues and abbreviations.

The phosphorylated serine residues are underlined in the sequence for clarity. The CTD sequence with 44 residues from the human CTD (residues between 1927 to 1970) is selected following the previous work^22, 29, 30^ which reported experimental NMR chemical shifts. CTD sequence with 14 residues contains two repeats of heptapeptide sequence. In addition, we explored 4 and 9 different phosphorylated states of CTD sequences with 44 and 14 residues, respectively.

Each of the CTD sequences on Table 1. were solvated in cubic boxes and when required, Na^+^ ions were included in order to neutralize systems. For the exp-CTD-non-phos system, we used both CHARMM C36m and CHARMM C36mw force fields (FFs), which the latter has a modification in non-bonded interactions between protein and water.^44^ We obtained a similar agreement with the experimental NMR chemical shifts^29, 30^ when using C36m and C36mw FFs (Figure 1 and Figure S1). C36mw provided more extended structures (Figure S2) that is expected for an IDP, therefore we used C36mw for the rest of the simulations. The TIP3P parameters^45^ were utilized for the explicit water. Then, an energy minimization was performed for 5000 steps with 100 kJ/mol tolerance. The systems were equilibrated for 625 ps while increasing the temperature from 100 K to 300 K. During the equilibration, the backbone and the side chains of CTDs were constrained using a force constant of 400 and 40 kJ/mol/nm^2^, respectively. The simulations were performed using OpenMM^46^ on GPU machines. Long-range electrostatic interactions were calculated using periodic boundary conditions with particle mesh Ewald (PME) algorithm.^47, 48^ The Lennard-Jones interactions were switched between 1.0 and 1.2 nm. The time step was set to 1 fs for the equilibration. Langevin thermostat was utilized with a friction constant of 1 ps^-1^ in order to maintain the temperature.

**Figure 1.**
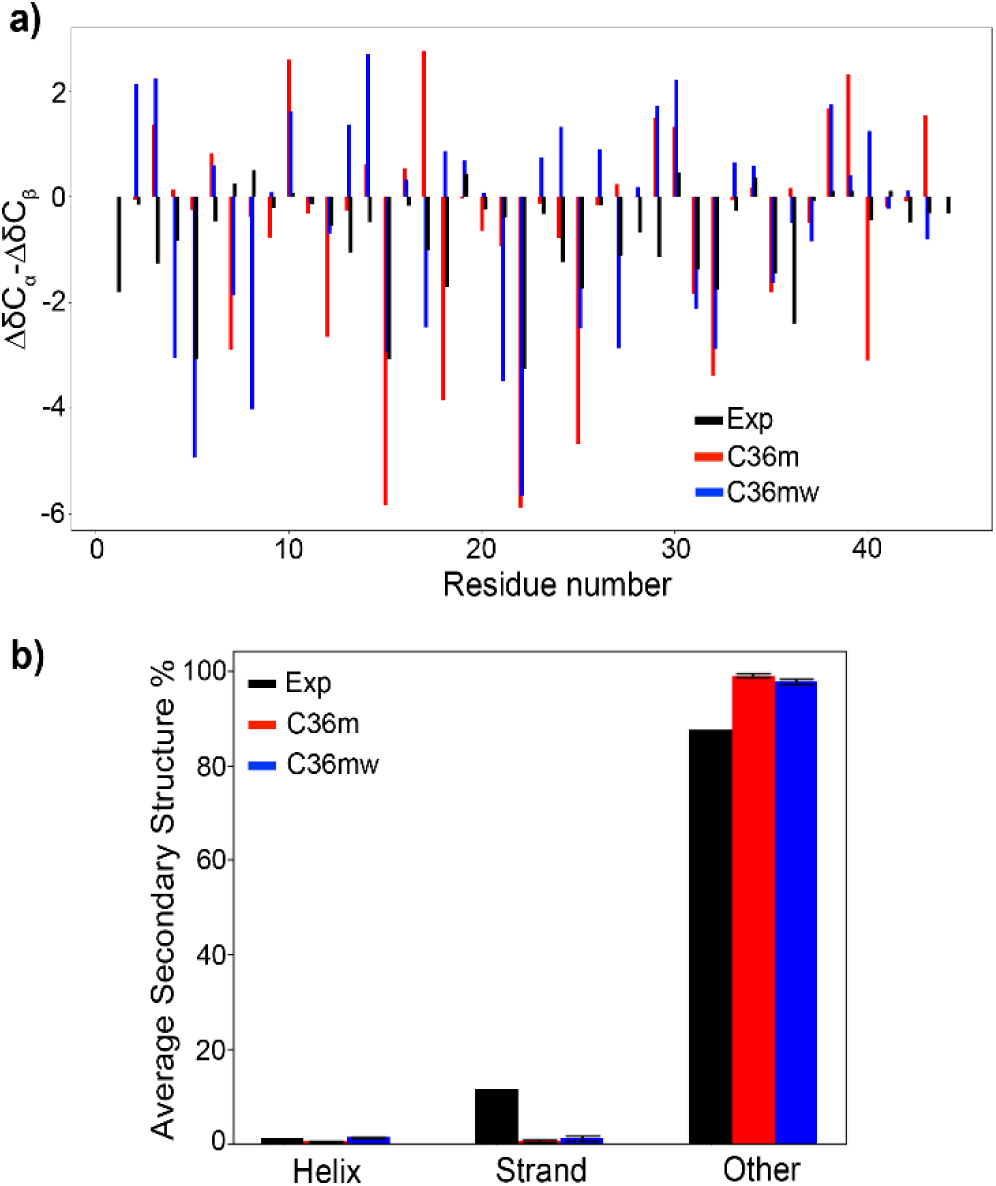
a) Comparison between the experimental secondary chemical shifts^29, 30^ (ΔδC_α_-ΔδC_β_) and the values derived from simulations using both C36m and C36mw FFs as a bar chart representation for exp-CTD-non-phos sequence, and b) average secondary structure predicted from NMR chemical shifts (experimental with δ2d software) and from simulations using C36m and C36mw FFs with DSSP program (error bars are calculated by splitting the full 200 ns trajectory in to 40 ns small trajectories).

### Replica-exchange molecular dynamics (REMD) simulations

All the production simulations were performed using REMD simulations in order to obtain an accurate conformational space for CTDs. REMD simulations were performed using OpenMM^46^ package with GPU enhanced environment. The final configurations from the previous equilibration step were utilized as the initial configurations for REMD simulations. For the CTD sequences with 44 residues and 14 residues on Table 1., 16 and 8 replicas were utilized respectively in order to maintain the exchange acceptance probability above 30 % between replicas (see Table S1). The temperature range for the replicas was set from 300 K to 500 K. Langevin dynamics was used as thermostat with the time step of 2 fs. Long-range electrostatic interactions were calculated using a reaction field approximation^49^ beyond a cutoff distance for the periodic systems. As for the equilibration, Lennard-Jones interactions were switched between 1.0 and 1.2 nm. At each 500 steps (time intervals of 1 ps) an exchange was attempted during all the REMD simulations. The production REMD runs were performed for 200 ns for each CTD system except for 2CTD-2P-5P-9P-12P shown in Table 1, which was extended up to 500 ns to obtain a better convergence compared to 200 ns trajectory (Figure S3). Frames were saved at every 10 ps during the simulations. Altogether, a total of 34.4 µs simulations were achieved.

### Data Analysis of the REMD simulations

The analyses were performed for the trajectories at the lowest temperature (300 K). The radius of gyration (R_g_), end-to-end distance, hydrogen bonds (H-bonds) analysis, distance maps and principal component analysis (PCA) were performed using MDAnalysis package.^50^ Distance maps were generated using the distances between center of masses of residues for each CTD. The PCA analysis was performed using the cartesian coordinates (backbone atoms) of the CTDs with first and second principal components (PC1 and PC2). The free energy landscapes (using PC1 and PC2 as well as R_g_ and end-to-end distance as reaction coordinates) and density distributions of R_g_ and H-bonds were generated by MATLAB.^51^ The weighted histograms in order to generate free energy landscapes were calculated using WHAM package developed by Grossfield lab.^52^ The secondary structures from the simulations were predicted using DSSP program^53^ in MDTraj.^54^ The secondary structures obtained from DSSP program were categorized as helix (alpha helix, 3/10 helix and pi helix), strand (isolated beta bridges and extended strands) and coil (loops, bends and turns). The experimental secondary structures were predicted using the δ2d software^55^ using available NMR chemical shifts for exp-CTD-non-phos system.^29, 30^ The δ2d software predicted α-helix, β-strand, coil and polyproline II structures, which the latter two are referred as other structures in this work. Chemical shifts from the simulations were calculated by SPARTA+ algorithm.^56^ Trajectories of full 200 ns were used for R_g_, end to end distance, PCA, H-bond calculations and secondary structure predictions, except for 2CTD-2P-5P-9P-12P which we used the last 400 ns of the full 500ns trajectory. Central structure was used for distance maps and chemical shift calculations. Central structure for each CTD was determined by calculating the average structure of CTD from the trajectory and then calculating the root-mean squared displacement (RMSD) between average structure and each frame of the trajectory. Then the frame with minimum RMSD with respect to the average structure was selected as the central structure. The Na^+^ ion densities around phosphate groups of serine residues were calculated using VolMap tool in the Visual molecular Dynamics package (VMD)^57^ (the Na^+^ densities were averaged along the trajectories and mapped as an iso-surface on the central structures of each CTD for clarity).

### Data Availability

An example of REMD simulation scripts, analysis scripts and most probable conformations obtained from the PCA analysis in PDB format were provided in a GitHub repository (https://github.com/bercemd/ctd_conformations).

## RESULTS

CTD of Pol II is a low complexity domain with a heptapeptide repeat, as human CTD has 52 repeats while yeast CTD has 26 repeats. It is computationally challenging to simulate the whole CTD from either human or yeast, while the model CTD with 44 residues, which was experimentally studied, and a CTD with 2 heptapeptide repeats (2CTD) could potentially provide important insights in the conformations of CTD sequences in general. We performed REMD simulations of the non-phosphorylated and phosphorylated 44-residue CTD and 2CTD models. Below, we first show the agreement of simulation results with the experimental NMR observations for the 44-residue CTD. Then, we present the results from the two CTD models in various phosphorylation states. Finally, we generalize our conclusion by proposing a model to explain phosphorylation effects on the conformations of CTD models.

### Simulations predicted mostly disordered conformations consistent with experiments

The secondary chemical shifts (ΔδC_α_-ΔδC_β_) calculated from NMR measurements^29, 30^ (experimental) and our simulations with two FFs (C36m and C36mw) are compared in Figure 1a for exp-CTD-non-phos sequence. Both C36m and C36mw showed varied agreements with the experimental secondary chemical shifts in Figure 1a, with 0.28 and 0.26 R^2^ correlation (Figure S1). For some residues both FFs presented a good agreement while there are deviations observed for most of the residues. However, chemical shift differences from the experiment and simulations with both FF suggest that there is not any significant helical propensity. Moreover, Figure 1b compares the average secondary structure percentages predicted from our simulations (using DSSP program) and from experimental NMR chemical shifts (using δ2d software). Average secondary structure percentages are in good agreement between experiments and simulations (both C36m and C36mw), specifically with helix and other structures. However, average strand structure percentage is higher from the experiments compared to two FFs which is consistent with a previous computational work with amber FF.^22^ Overall, both FFs provided a mostly disordered conformation that is in good agreement with the experimental results.

### Phosphorylation caused extended conformations in CTD sequences with 44 residues

We performed REMD simulations of 44-residue CTD (exp-CTD-non-phos) and its four phosphorylated structures (exp-CTD-5P-40P, exp-CTD-5P-22P-40P, exp-CTD-5P-12P-18P-32P and exp-CTD-5P-12P-18P-25P-32P-40P). In order to obtain the most probable low energy conformations and the conformational space of each CTD sequence, we applied principal component analysis (PCA) using cartesian coordinates. Figure 2 shows the free energy landscapes generated by PCA analysis using the first and second principal components (PC1 and PC2 respectively) and the lowest energy conformations in the bottom (X_1_-X_7_). In all cases, the plots show rugged energy landscapes and a wide range of conformational space for different CTD sequences which is expected for IDPs. Conformations of phosphorylated CTDs (X_2_ to X_7_) were more extended compared to the non-phosphorylated CTD (X_1_). In addition to this, Figure S4 shows the free energy landscapes for R_g_ vs. end-to-end distances which suggest that more extended structures were observed upon phosphorylation, regardless of the number and position of the phosphorylation sites.

**Figure 2.**
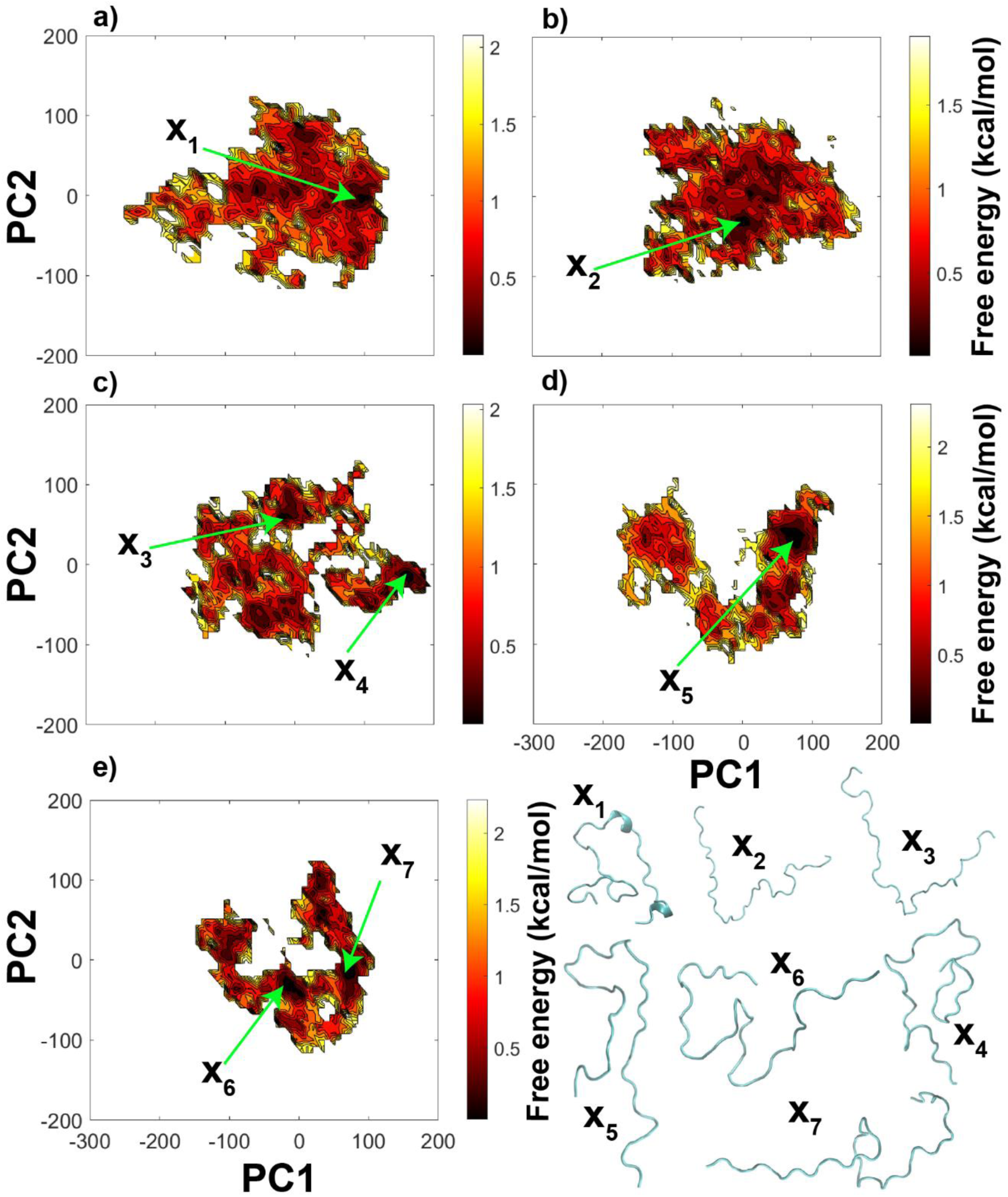
The free energy landscapes using PC1 and PC2 as reaction coordinates from the principal components analysis (PCA) using cartesian coordinates of CTDs, a) exp-CTD-non-phos, b) exp-CTD-5P-40P, c) exp-CTD-5P-22P-40P, d) exp-CTD-5P-12P-18P-32P and e) exp-CTD-5P-12P-18P-25P-32P-40P. In addition, X_1_-X_7_ represents a few of the lowest energy conformations of different CTD sequences.

Figure 3a shows the R_g_ density distributions for CTD sequences with 44 residues. All the phosphorylated states of the CTD show expansion with respect to the non-phosphorylated state (exp-CTD-non-phos). This observation is expected as the net charge of exp-CTD-non-phos sequence is negative and upon phosphorylation the repulsive interactions between negatively charged residues and the phosphate groups tend to be increased and consequently the conformations were extended. This observation is also in agreement with the previous computational work done by Jin & Gräter^58^ that they observed extended structures for the IDP sequences with negative net charge and around same length as 44-residue CTD sequences upon phosphorylation. In addition, compared to other phosphorylated states exp-CTD-5P-22P-40P sequence shows a broader density distribution of R_g_, while exp-CTD-5P-12P-18P-25P-32P-40P has sharper distribution. This suggests that the relative positions of the phosphorylated residues might play an important role. There is an opportunity to sample more diverse conformations for exp-CTD-5P-22P-40P sequence with well spread relative positions of the phosphorylated residues. However, an increased number of phosphorylation sites restricted conformational sampling in the case of the exp-CTD-5P-12P-18P-25P-32P-40P model.

**Figure 3.**
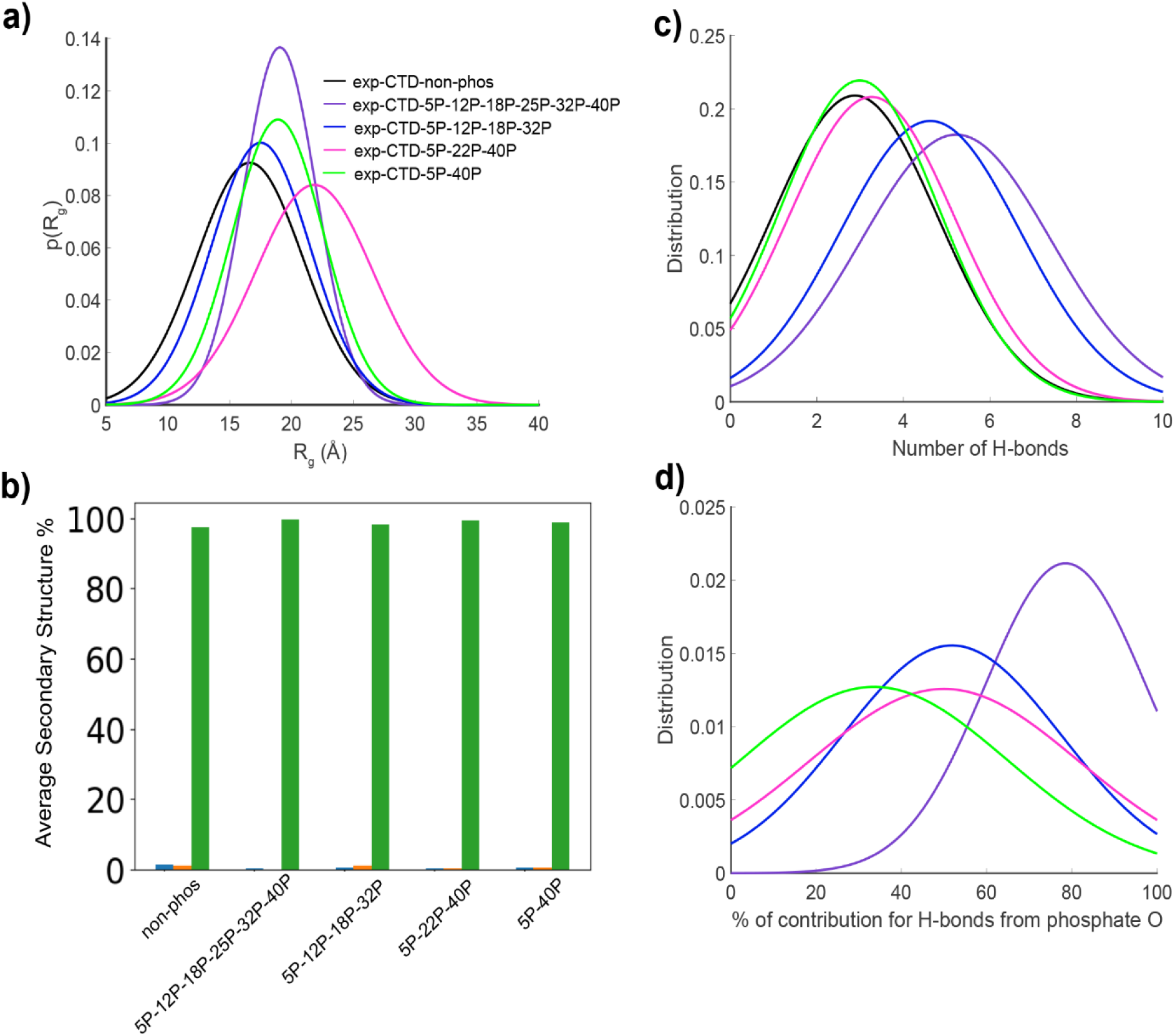
a) The radius of gyration (R_g_) density distributions for CTD sequences with 44 residues, b) average secondary structure percentages for all the CTD systems with 44 residues as a bar chart representation (blue – helix, orange - strand and green – coil), c) distributions of total number of intrapeptide H-bonds for CTD systems with 44 residues and d) distributions of contributions from oxygen atoms of phosphate groups to intrapeptide H-bonds of phosphorylated CTDs with 44 residues. (Colors of the distribution curves are same as panel a) for panel c) and d)).

Figure 3b represents the overall average secondary structure for the CTDs with 44 residues and Figure S5 shows the percentages along each residue of the CTD sequence. As expected, all the CTDs are mostly disordered which can be verified with high percentages (>90%) of coil structures. Here coil structures include all the loops, bends and turns according to DSSP definition.^53^ Also, Figure 3b and Figure S5 show helix structures (alpha helix, 3/10 helix and pi helix) and strand structures (isolated beta bridges and extended strands) in lower percentages (<10%). We also generated the secondary structure evolution with time in Figure S6 which helped us to understand the stability of these secondary structures of low percentages in Figure 3b (see Figure S5 also). We found that both helix and strand structures remain stable for very short periods of time, compared to where in most cases coil structures remain stable for longer periods of time. Moreover, exp-CTD-non-phos shows helix and strand structures for almost all the residues in low percentages and more elevated helix structures around residue IDs 24-32 and 36-44 (see Figure S5a and Figure S6a) compared to the phosphorylated CTDs, which can be also seen in the minimum energy conformation obtained by PCA (see X_1_ conformation in Figure 2). A previous study by Tang et al.^22^ also reported mostly disordered structure with low helix probabilities for the same non-phosphorylated CTD sequence using an AMBER force field. Phosphorylation did not show any significant change in the secondary structures, although there are slightly more elevated strand structures for exp-CTD-5P-12P-18P-32P (see Figure S5d) around residue IDs 6-14 and 27-30. Overall, CTDs with or without phosphorylation show high coil structure percentages and low helix and strand percentages, however, there are few specific changes of secondary structure in low percentages upon phosphorylation.

In order to identify the interactions with nearby residues in CTD conformations, we analyzed intramolecular H-bonds. Figure 3c and 3d show the distributions of the total number of intrapeptide H-bonds for CTDs with 44 residues and the contributions from the oxygens of phosphate groups to the total number of H-bonds, respectively. We found that the total number of intrapeptide H-bonds formed between residues of CTD increases with the number of phosphorylated residues. Consistently, the contribution to the total number of intrapeptide H-bonds from oxygens of phosphate groups also increases with the number of phosphate groups. In addition to this, Figure S7 shows that the number of close contacts increases around the phosphorylated residues according to the distance maps for exp-CTD-5P-12P-18P-32P and exp-CTD-5P-12P-18P-25P-32P-40P. This shows that the number of overall interactions increases around the phosphorylation sites in the CTDs for highly phosphorylated systems.

### Phosphorylation had diverse effects on the conformation of CTD sequences with 14 residues

The CTD model with 44 residues showed a broad conformation landscape and increased extension in the structure upon phosphorylation. In order to cover effects of a larger set of phosphorylation patterns to the conformation we studied a shorter CTD model, which has 2 heptapeptide repeats (2CTD). Figure 4 represents the free energy landscapes from the principal component analysis and a few of the low energy conformations for each 2CTD model. All the free energy landscapes are very broad and mostly exhibit a large conformational space. In addition, local energy minimal regions are also broad and separated by low energy barriers compared to the CTDs with 44 residues. This verifies that 2CTDs have many low energy conformations which can interchange within each other. The structures in Figure 4 show that 2CTD-2P-5P-9P-12P, 2CTD-2P-5P-12P, 2CTD-2P-12P and 2CTD-5P-12P have relatively more contracted low energy conformations that are Z_5_, Z_6_, Z_8_ and Z_10_ respectively (also see A_5_, A_6_, A_8_ and A_10_ conformations in Figure S8 in the SI).

**Figure 4.**
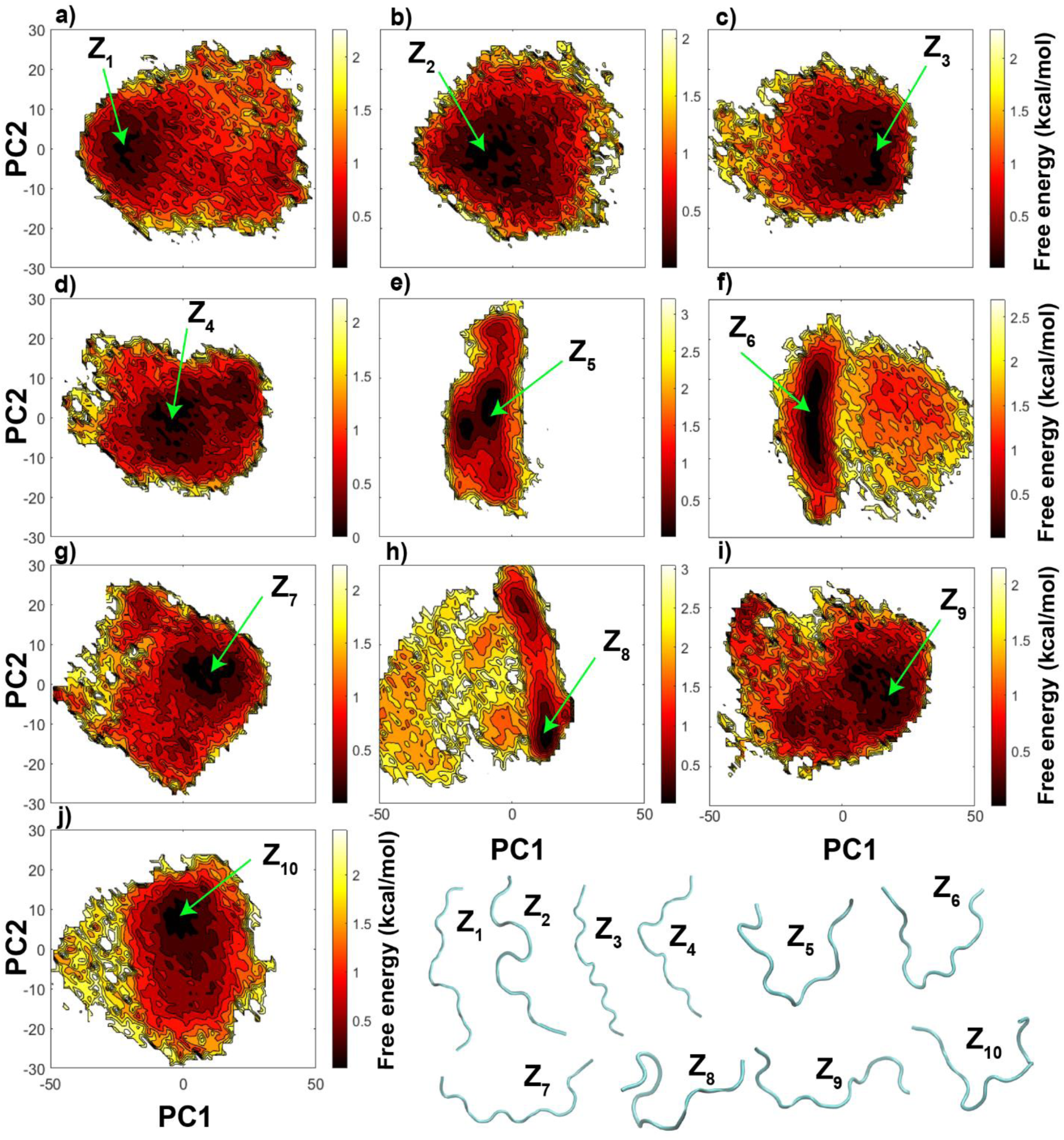
The free energy landscapes using PC1 and PC2 as reaction coordinates from the principal components analysis (PCA) using cartesian coordinates of CTDs, a) 2CTD-non-phos, b) 2CTD-2P, c) 2CTD-2P-5P, d) 2CTD-2P-5P-9P, e) 2CTD-2P-5P-9P-12P, f) 2CTD-2P-5P-12P, g) 2CTD-2P-9P, h) 2CTD-2P-12P, i) 2CTD-5P and j) 2CTD-5P-12P. In addition, Z_1_-Z_10_ represents a few of the lowest energy conformations of different CTDs.

Figure 5a demonstrates the density distributions of R_g_ for CTDs with 14 residues. All the density distributions of R_g_ are very broad for CTDs with 14 residues except for 2CTD-2P-5P-9P-12P suggesting that the conformations for most of the 2CTDs are interconverting within a large conformational space. Phosphorylation in shortened CTD results in complex changes in R_g_. 2CTD-2P-5P, 2CTD-2P-5P-9P and 2CTD-2P-9P show expansion upon phosphorylation, while a few of the 2CTDs show contraction mainly the sequences 2CTD-2P-5P-9P-12P, 2CTD-2P-5P-12P, 2CTD-2P-12P and 2CTD-5P-12P. These observations suggest that for the CTD with a shortened length (compared to 44 residues), in addition to the net charge and the number of phosphorylated residues, the relative positions of phosphorylation sites also determine whether the peptide will expand or contract. Interaction with nearby residues and ion coordination might be some of the factors that are contributing to the changes in conformational space upon phosphorylation. Similar type of observation was discussed in a previous computational study by Rieloff & Skepö for IDPs around similar lengths.^23^ In that study, they observed an expansion for Tau1 (IDP with positively charged non-phosphorylated state and 11 residues) and a contraction for β-casein (IDP with negatively charged non-phosphorylated state and 25 residues) after phosphorylation.

**Figure 5.**
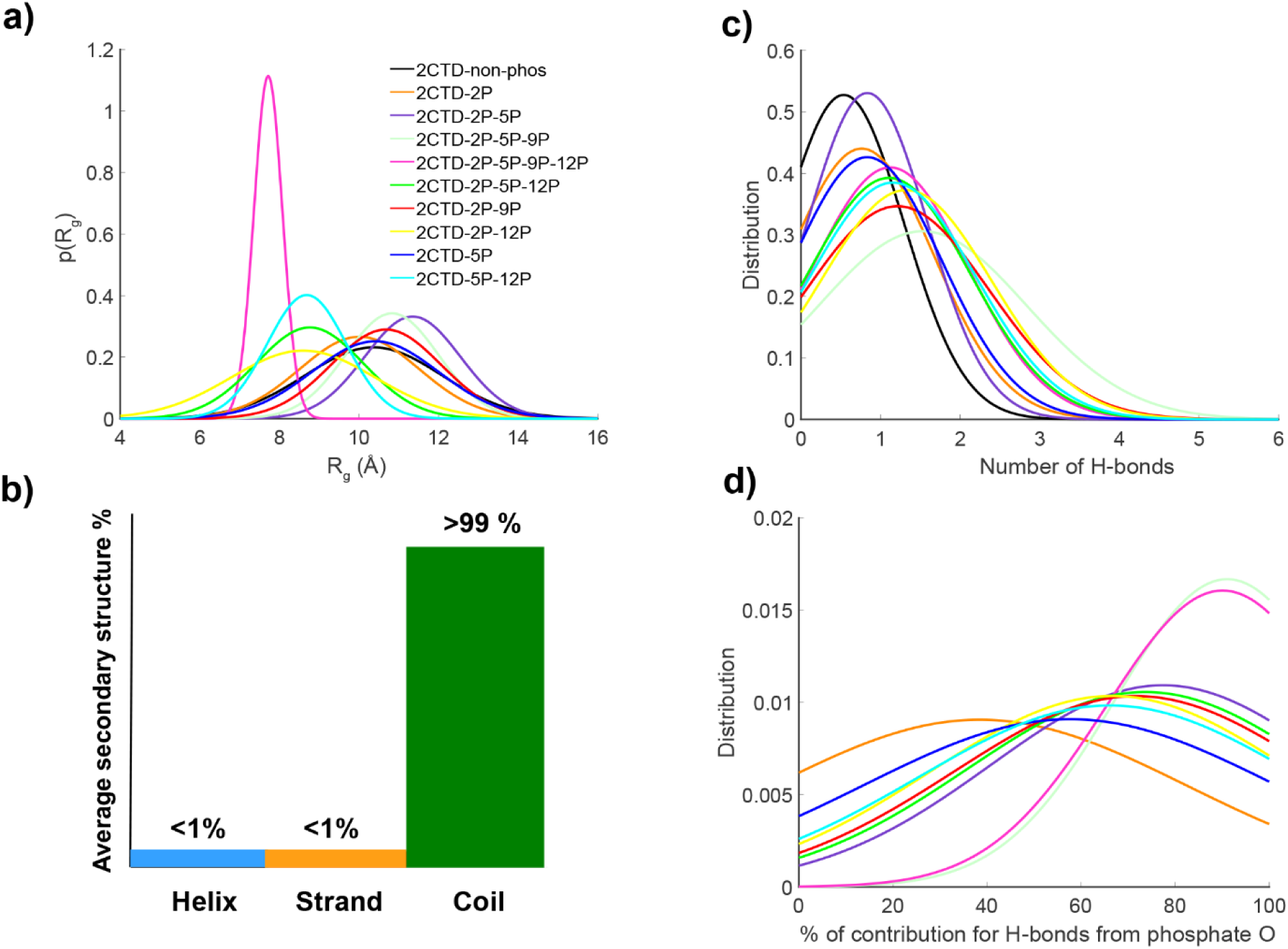
a) The radius of gyration (R_g_) density distributions for CTD sequences with 14 residues, b) average secondary structure percentages for all the 2CTD systems as a common bar chart, c) distributions of total number of intrapeptide H-bonds for 2CTD systems and d) distributions of contributions from oxygen atoms of phosphate groups to intrapeptide H-bonds of phosphorylated 2CTD systems. (Colors of the distribution curves are same as panel a) for panel c) and d)).

The large conformational spaces observed in CTD with 14 residues can be verified further from the average secondary structure percentages in Figure 5b. For all the 2CTDs, average coil structure (loops, bends and turns) percentage is more than 99% and average helix (alpha helix, 3/10 helix and pi helix) and strand (isolated beta bridge and extended strand) percentages are less than 1 % for the entire protein (Figure 5b) and for every residue (Figure S9), which suggests that all the 2CTDs are more disordered than the CTDs with 44 residues (>90% average coil structure). Even for a few of the 2CTDs average coil structure percentage is 100% (see Figure S9c-e). Also, the helix and strand structures predicted for 2CTDs are only stable for very short periods of time as shown in Figure S10.

In order to understand the interactions behind the conformational changes in 2CTD upon phosphorylation, we analyzed intrapeptide H-bonds. Figure 5c shows that there is an increase of the total number of intrapeptide H-bonds upon phosphorylation of 2CTDs as we observed for CTDs with 44 residues. In some cases, the increase of H-bonds is not significant (2CTD-2P, 2CTD-2P-5P and 2CTD-5P in Table S2), while, in other cases the increase was more than two-fold (see 2CTD-2P-5P-9P, 2CTD-2P-5P-9P-12P, 2CTD-2P-5P-12P, 2CTD-2P-9P, 2CTD-2P-12P and 2CTD-5P-12P in Table S2). We observed a similar pattern of increments in CTDs with 44 residues in Table S2. This means in some cases the phosphorylation significantly induces the number of intrapeptide H-bonds formed in CTDs with both 44 and 14 residues which will eventually determine their conformations. Also, as we observed for CTDs with 44 residues, the contribution to the total number of intrapeptide H-bonds is mostly from the oxygens of phosphate groups for 2CTD systems (see Figure 5d). Distance maps in Figure S11 show that some of the contacts were lost upon phosphorylation while there are few close contacts that appeared and specifically linked with the phosphorylated residues. These close contacts appear mostly in 2CTD-2P-5P-9P-12P, 2CTD-2P-5P-12P, 2CTD-2P-12P and 2CTD-5P-12P which are the 2CTDs with more contractions compared to 2CTD-non-phos (non-phosphorylated state) in Figure 5a.

In order to understand the electrostatic interactions that potentially stabilize the contracted conformations upon phosphorylation, we analyzed Na^+^ ion densities. Figure 6 shows the average Na^+^ ion density around phosphate groups of the central structures of contracted 2CTDs, which are 2CTD-2P-12P, 2CTD-2P-5P-12P, 2CTD-5P-12P and 2CTD-2P-5P-9P-12P. This figure demonstrates Na^+^ ion coordination by the negatively charged phosphorylated Ser residues and bending of the structure as a result of this coordination. This result provides an explanation for the decreased R_g_ values and increased contractions of the conformations observed in these specific 2CTD phosphorylated models shown in Figure 5a. As we visualize the low energy conformations specifically for 2CTD-2P-5P-9P-12P, 2CTD-5P-12P, 2CTD-2P-12P and 2CTD-2P-5P-12P, interactions with Na^+^ ion makes the phosphorylated residues come closer to each other, which would be energetically unfavorable otherwise due to the presence of repulsive forces as the phosphate groups are negatively charged. Compared to the 44-residue CTDs, due to the shortened length of 2CTDs, Na^+^ ions can form stable complexes with phosphate groups through electrostatic interactions, when phosphate groups are especially present near the two terminal ends of 2CTDs. All the panels in Figure 6 show high Na^+^ ion density around phosphate groups which eventually induce more compact conformations for the above mentioned 2CTDs. In contrast, the other 2CTD models show more widely distributed Na^+^ ion densities that did not support any bending and contraction in the structures (see Figure S12). The bottom-line is that the shortened length of 2CTDs combined with the spacings between multiple phosphorylated sites allow some of the 2CTDs to have more contracted conformations with help of the formation of Na^+^-phosphate group complexes compared to the non-phosphorylated state even though the net charges of those phosphorylated 2CTDs are negative.

**Figure 6.**
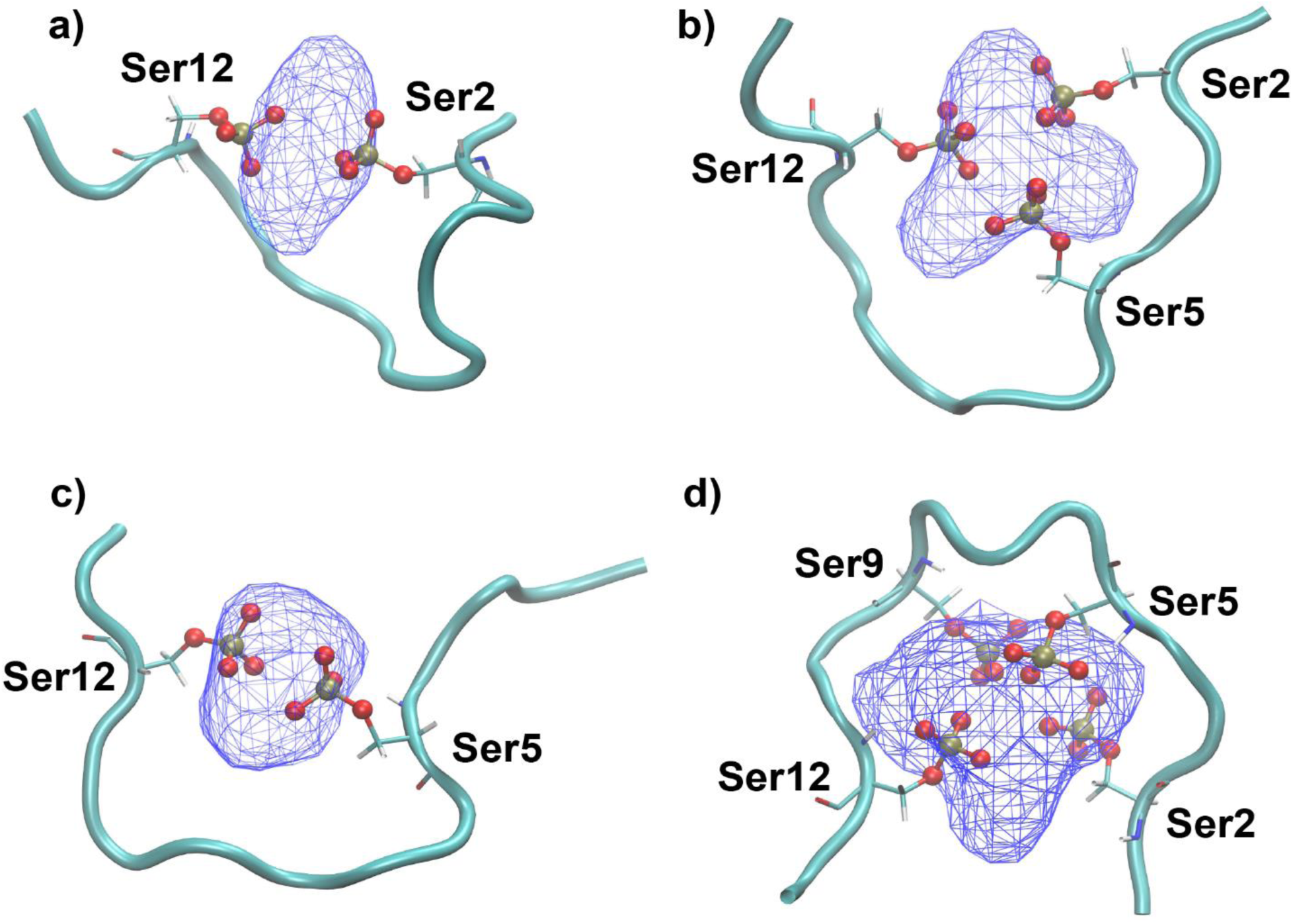
The average Na^+^ ion density around the phosphate groups in serine (Ser) residues for the central structures of contracted 2CTDs. a) 2CTD-2P-12P, b) 2CTD-2P-5P-12P, c) 2CTD-5P-12P and d) 2CTD-2P-5P-9P-12P. Blue wire mesh represents the density of Na^+^ ions around the phosphate groups of highlighted serine residues of each 2CTD sequence. The cyan cartoon structure shows the backbone of each 2CTD sequence. Red and yellow spheres represent oxygen and phosphorus atoms respectively. The visualizations were generated using visual molecular dynamics (VMD) package.^57^

### Length of CTD and relative positions of phosphorylation sites affect the conformations

We observed that phosphorylation of CTD with 44 residues caused extended structures, while CTD with 14 residues have either extended or contracted structures upon phosphorylation. A simple model shown in Figure 7 suggests an explanation of the distinct effects of phosphorylation. As the ion density analysis showed, the 2CTD structures bend when multiple phosphorylation sites are in a certain distance range (7-10 residues apart) to coordinate Na^+^ ion by negatively charged oxygen atoms. Bending of the structures causes more compact conformations for such cases. In contrast, for the 44 residue CTDs, phosphorylated structures tend to be extended, and bending is not supported potentially due to the high entropic cost to bend longer disordered structures. We also note that, for the 2CTDs, not only distance between the phosphorylation sites, but also their relative locations affect the compactness of the structure. For example, the effects of phosphorylation for 2CTD-5P-12P and 2CTD-2P-9P are different, although their phosphorylation sites are both 7 residues apart (Figure 6). Ion coordination by two phosphorylated serine residues of 2CTD-5P-12P supported bending, while in the 2CTD-2P-9P system, the phosphorylated serine residues at position 2 and 9 were coordinating Na^+^ ions separately (Figure S12). In this case, the hydrogen bonds between phosphorus oxygens at position 9 and the neutral serine residue at position 7 stabilize the structure and prevent bending. Overall, our model suggests that ion coordination can cause contraction in the short CTD structures, whereas for the longer structures, contraction was restricted due to the combination of electrostatic repulsion and the entropic cost for bending.

**Figure 7.**
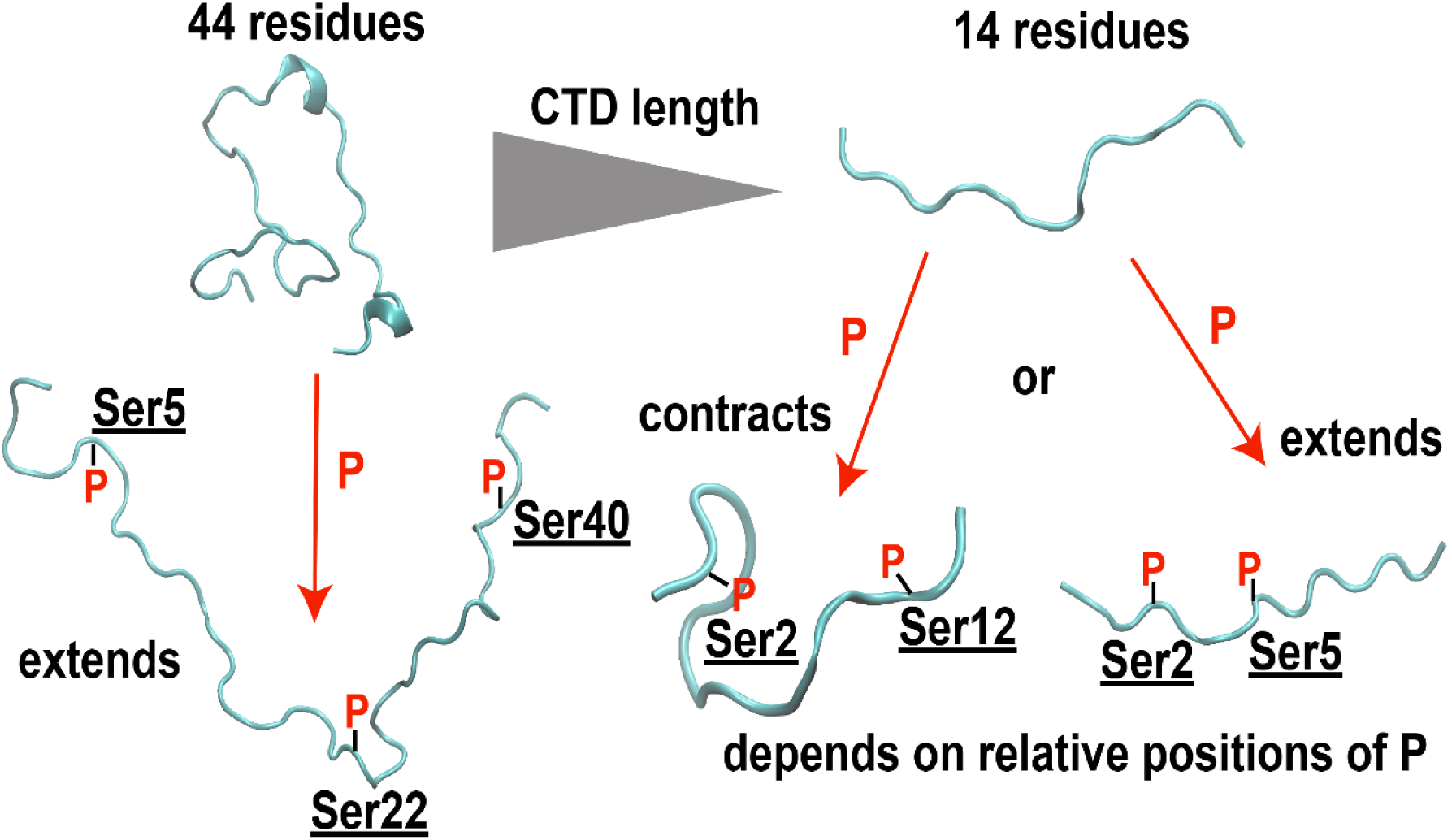
Model for the effects of phosphorylation in conformations of the CTDs at different lengths and phosphorylation patterns.

## DISCUSSION

In this study, we generated conformational ensembles of CTD models at two different lengths and varying phosphorylation states using enhanced sampling MD simulations. REMD simulations at all-atom details provided a large span of conformational ensemble for both 14- and 44-residue CTD models. As secondary structure predictions of 44-residue CTD show (Figure 1), the five blocks of simulations provided small standard errors that suggest the convergence of the simulations. Additionally, secondary structures evolvement through time (Figure S6 and S10) suggests that coil structures are predominant while helix and beta structures are formed transiently, which also support the convergence of secondary structures for the simulations. However, the computational cost for CTD models substantially increased for the 44-residue CTD as we needed to run 16 replicates to cover the most probable conformational spaces. Therefore, generating the full span of conformational space for the yeast or human CTDs, which have 26 and 52 heptapeptide repeats respectively, is computationally challenging using atomistic REMD simulations. Alternative strategies can be applied for studying full length CTDs, which include coarse-grained MD simulations,^16, 59–61^ fragmentation of long chains^13, 14^ or generative machine learning models^19–21^ using variational auto-encoder or attention-based approaches. A future direction for our study is to generate conformations of longer CTD models with different phosphorylation patterns using coarse-grained simulations and then develop a machine learning model based on coarse-grained conformations to predict conformational ensembles of CTD sequences in any given length and phosphorylation pattern. Similar studies showed that such approaches successfully predict the conformations,^19–21^ while none of these studies include post-translational modifications.

CTD of Pol II is known to undergo several post-translational modifications, including phosphorylation of serine residues at the 2^nd^, 5^th^ and 7^th^ positions of the heptapeptide repeat. There is a large number of possible combinations of phosphorylation within full length CTDs and it is not entirely known which phosphorylation patterns can take place together or which ones are mutually exclusive *in vivo*.^26, 62^ In our study, we selected all the possible 2Ser and 5Ser combinations from the N- to C-terminal ends for 2CTD model as the 2Ser and 5Ser positions are known to be the most observed phosphorylation sites for CTD.^63, 64^ However, we note that some of the phosphorylation states we explored in this study may not take place *in vivo*, and, therefore, may be biologically irrelevant. Although the CTD phosphorylation pattern is not entirely clear, some studies suggest that mono-phosphorylation is more common and adjacent phosphorylation increases the prevalence of double phosphorylation in a repeat;^63^ and phosphorylation level is distributed evenly across the sequence that similar amount of phosphorylation is observed close to the Pol II core and at the end of the tail.^64^ Phosphorylation pattern may also be related to the steric hindrance or accessibility of the positions for the kinases which are the proteins that catalyze phosphorylation. Regardless of biological feasibility of the phosphorylation patterns, we explored a large set of potential positions for the 2CTD to obtain some generalized rules for the effects of phosphorylation on the conformations.

We proposed a model to describe the effects of phosphorylation on the conformation of CTDs at different lengths. Phosphorylation introduces an increased electrostatic repulsion, which causes an extension of the structure in a longer CTD (44-residue) as expected, while the effects are more complicated for a relatively shorter CTD (14-residues) as the repulsive interactions can be compensated by Na^+^ ion coordination. We concluded that the contraction of the structures is allowed in the 2CTDs but not in the 44-residue CTDs as the entropic cost for bending is relatively smaller for 2CTD. However, if we go to even longer sequences, the conformational space will be larger and there could be more complicated alterations in conformations at different phosphorylation patterns. But, overall, we expect that there would be even higher entropic barriers for a significant bending for the longer CTDs upon phosphorylation and consequently more extended structures can form as was reported earlier.^33^ Furthermore, our model does not include other factors that can affect conformations, including ion concentrations, binding of CTD to other proteins and crowding of the cellular environment. We performed the simulations at neutral charge that only counter ions were present in the system. We would expect that a higher salt concentration may decrease extension of CTDs at longer lengths due to the screening of the electrostatic interactions. Additionally, CTD of Pol II is known for interacting with other proteins such as mediator complex, capping enzymes, transcription and elongation factors.^26, 65^ The conformation of CTD may alter upon binding, which is not addressed by our model. As a last point, highly concentrated cellular systems may also induce relatively more compact structures for CTDs and may force intermolecular CTD interactions that may alter the conformations as well.

CTD of Pol II is also well recognized for its involvement in LLPS formation^27, 28^ and phosphorylation of CTD was suggested to regulate such phase separation events.^28, 66^ Therefore, it is crucial to determine conformational changes upon phosphorylation to better understand its effects on phase separation. To investigate phase separation by CTD, coarse-grained models in conjunction with enhanced simulation techniques can be applied as was done extensively in recent studies for similar systems.^67–69^ However, available coarse-grained models may need to be fine-tuned for CTD sequences and especially for the phosphorylated serine residues. One way to do this is to parameterize a coarse-grained model against all-atom MD simulations of concentrated CTD systems, which will be one of our future research directions.

## CONCLUSIONS

We reported computationally generated conformational ensembles of Pol II CTD at different lengths and phosphorylation states. We predicted highly disordered structures for all the systems with predictions of less than 10% of helix and beta strand structures. Introduction of phosphate groups on the serine residues caused more extended structures for the long CTD, while contraction of structure is observed for some of the short CTD systems. We proposed a model that summarizes the effects of phosphorylation on the conformation of the CTD systems. According to our model, Na^+^ ion coordination by multiple phosphate groups takes place depending on the relative positions of the phosphorylation sites and it stabilizes bending structures for short CTDs that causes contraction. On the other hand, long CTD extends its structure upon phosphorylation due to the increased electrostatic repulsion and entropic cost for bending. Future studies will focus on simulating CTD models in concentrated systems to fine-tune coarse-grained models, which will be later used to obtain conformations of full-length CTDs and to study LLPS formation by CTD interactions.

## ASSOCIATED CONTENT

### Supporting Information

The supporting information is available free of charge.

Linear regression analysis for secondary chemical shifts from experiments and simulations; comparison of free energy landscapes using R_g_ and end to end distance as reaction coordinates between C36m and C36mw FFs for exp-CTD-non-phos; table of total acceptance ratios for 2CTD and CTDs with 44 residues for REMD simulations; comparison of PCA profiles for 2CTD-2P-5P-9P-12P from 200 ns and 500 ns trajectories; free energy landscapes using R_g_ and end to end distance as reaction coordinates for CTDs with 44 and 14 residues; secondary structure time evolutions and average secondary structure % along sequences for CTDs with 44 and 14 residues; contact maps generated using distances between center of masses of residues for CTDs with 44 and 14 residues; table of average number of intrapeptide H-bonds for CTDs with 44 and 14 residues and average Na^+^ densities around phosphate groups of other 2CTDs compared to Figure 6 in main text.

## AUTHOR INFORMATION

### Corresponding Author

Bercem Dutagaci − Department of Molecular and Cell Biology, University of California, Merced, Merced, California 95343, United States.

### Authors

Weththasinghage D. Amith - Department of Molecular and Cell Biology, University of California, Merced, Merced, California 95343, United States.

### Notes

The authors declare no competing financial interest.

## ACKNOWLEDGMENTS

The authors used the computational resources at the MERCED cluster and PINNACLES cluster at the University of California Merced and at the National Science Foundation’s Extreme Science and Engineering Discovery Environment (XSEDE) facilities now known as Advanced Cyberinfrastructure Coordination Ecosystem: Services & Support (ACCESS) under the grant TG-BIO210145. We also thank Prof. Nicolas L. Fawzi for providing information about experimental NMR chemical shifts data for 44-residue CTD.

## TOC Entry

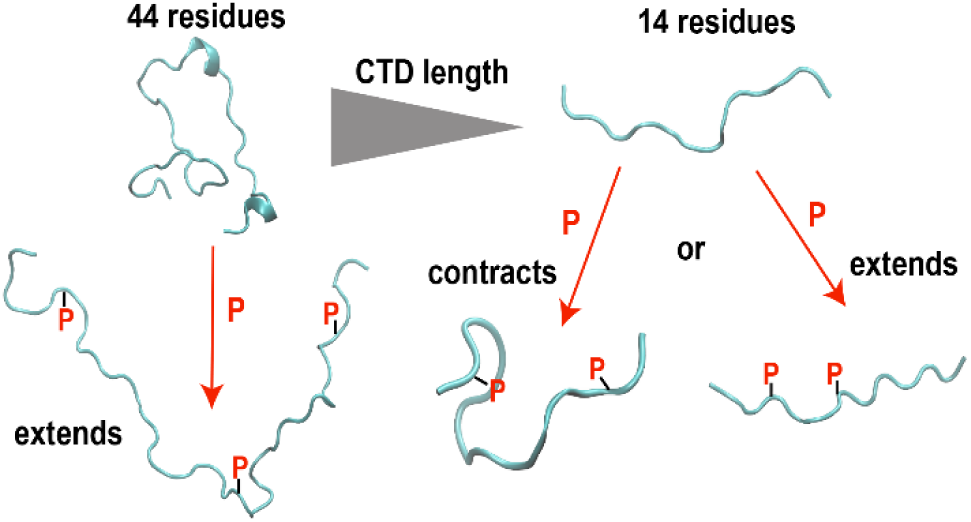

