## Supporting Information for "Complex Conformational Space of RNA Polymerase II C-Terminal Domain upon Phosphorylation"

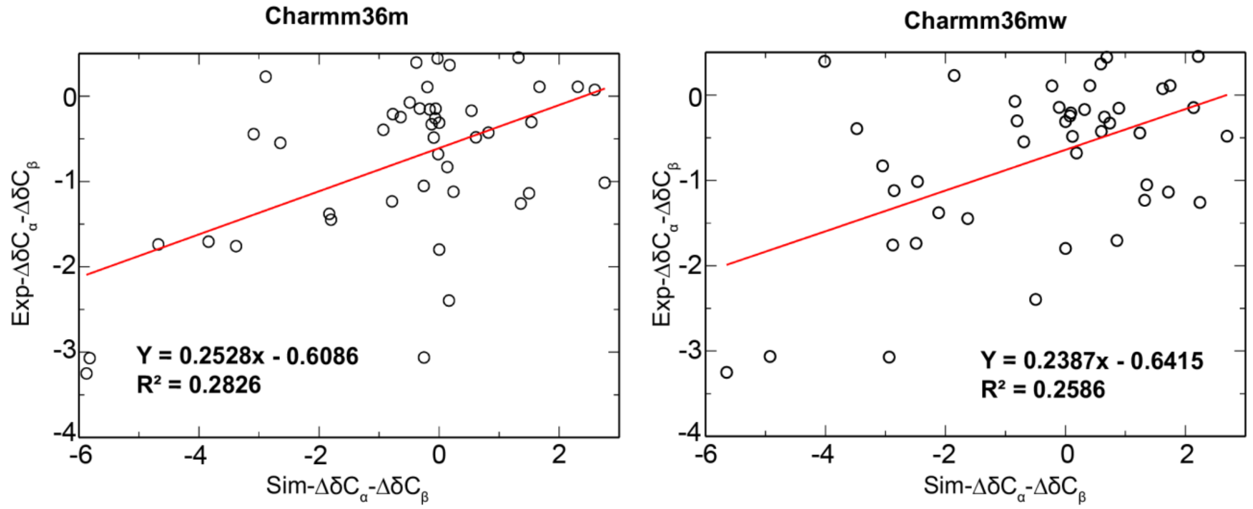


**Figure S1:** Comparison between the experimental secondary chemical shifts ($\Delta\delta C_\alpha$-$\Delta\delta C_\beta$) and secondary chemical shifts determined from C36m and C36mw FFs for exp-CTD-non-phos sequence with linear regression analysis (trendlines are shown using red solid lines and the equations for the trendlines and $R^2$ values are displayed on the plots).

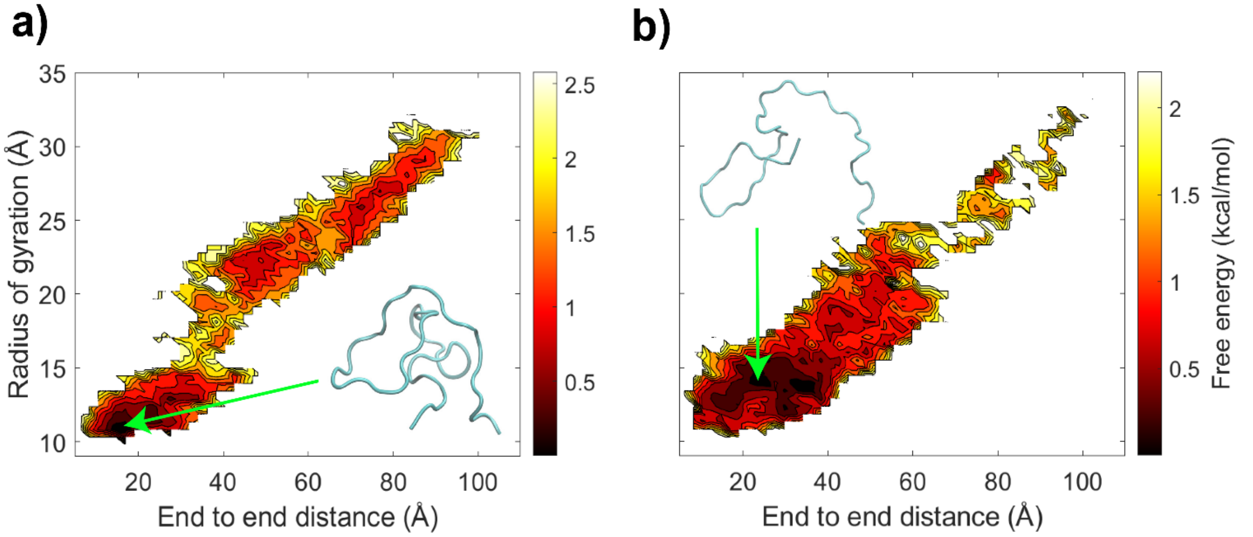


**Figure S2:** The free energy landscape using radius of gyration and end to end distance as reaction coordinates for exp-CTD-non-phos with 44 residues from the simulations with C36m (a) and C36mw (b) force fields. Low energy conformations are shown in cartoon representations.

**Table S1:** Total acceptance ratios between the neighboring replicates for REMD simulations of CTD models with 14 (8 replicates) and 44 (16 replicates) residues.

| System | Acceptance ratio (%) |
|---|---|
| exp-CTD-non-phos | 34.774 |
| exp-CTD-5P-40P | 34.772 |
| exp-CTD-5P-22P-40P | 34.777 |
| exp-CTD-5P-12P-18P-32P | 34.783 |
| exp-CTD-5P-12P-18P-25P-32P-40P | 34.783 |
| 2CTD-non-phos | 36.352 |
| 2CTD-2P | 36.377 |
| 2CTD-2P-5P | 36.348 |
| 2CTD-2P-5P-9P | 36.374 |
| 2CTD-2P-5P-9P-12P | 36.366 |
| 2CTD-2P-5P-12P | 36.361 |
| 2CTD-2P-9P | 36.413 |
| 2CTD-2P-12P | 36.359 |
| 2CTD-5P | 36.364 |
| 2CTD-5P-12P | 36.379 |

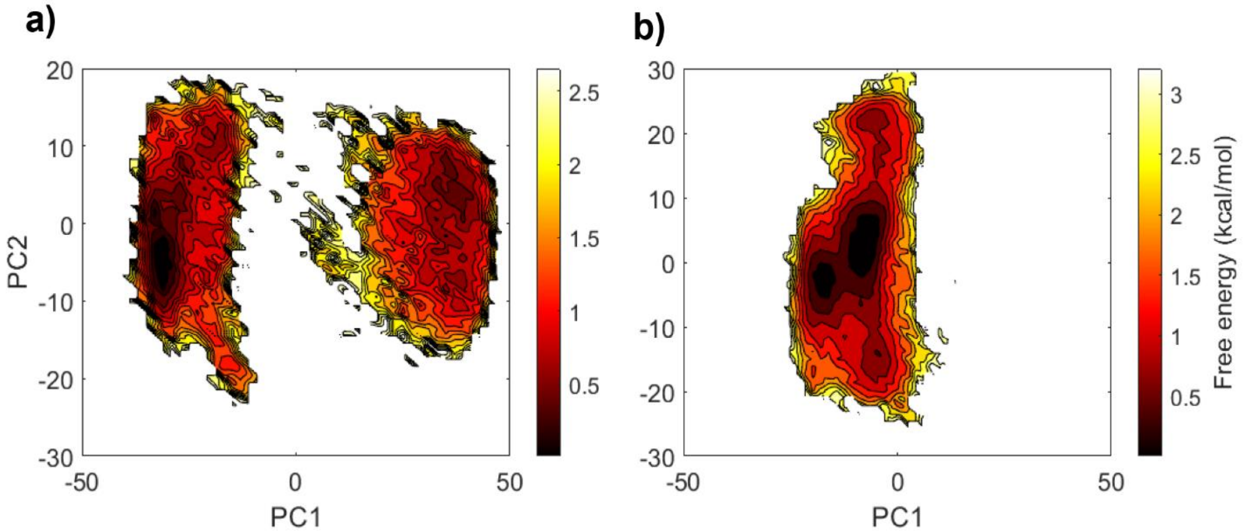


**Figure S3:** The free energy landscapes from PCA analysis for 2CTD-2P-5P-9P-12P system using first 200 ns of the simulation (a) and the 500 ns simulation (last 400 ns used for plot) (b).

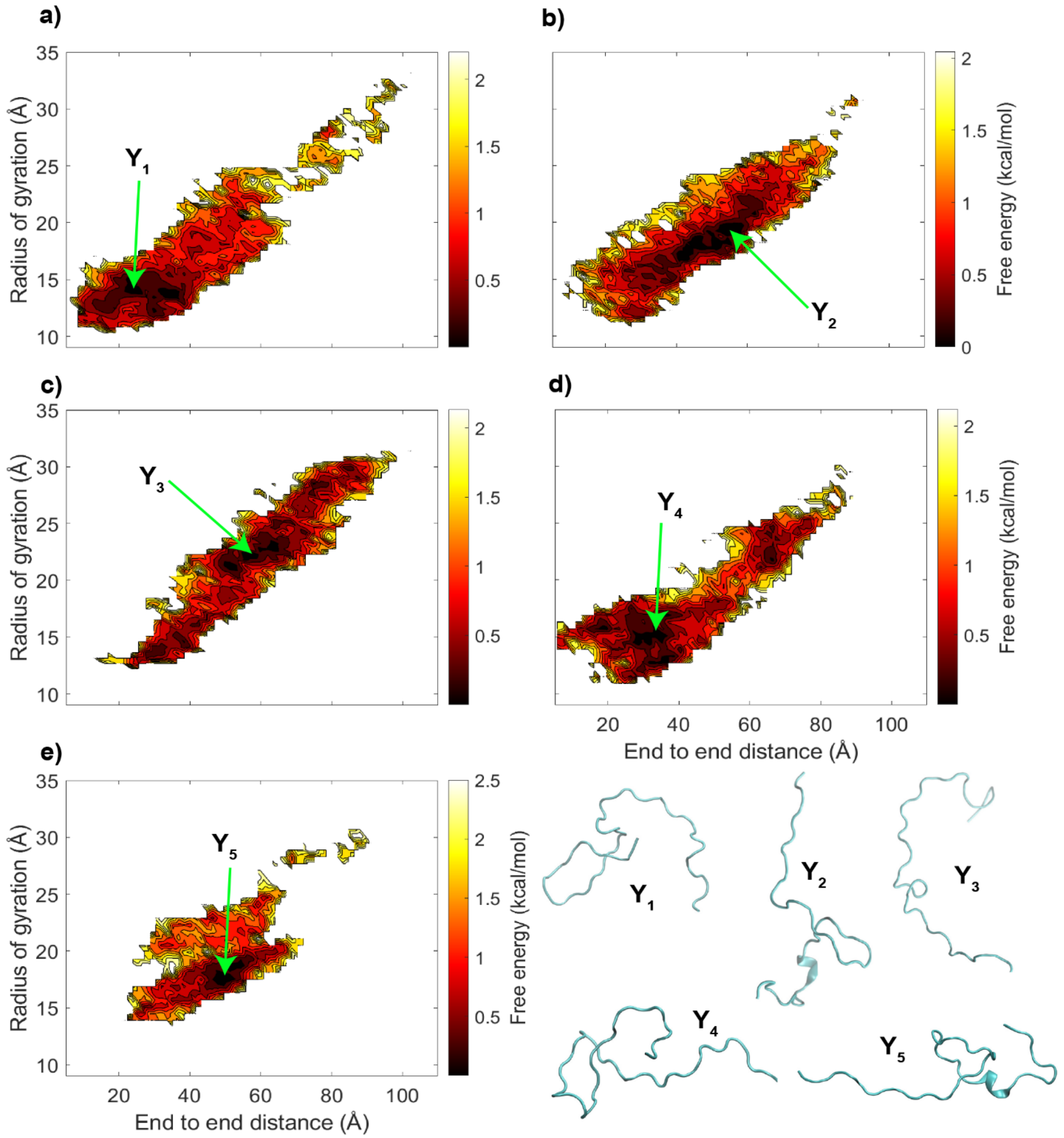


**Figure S4:** The free energy landscapes using radius of gyration ($R_g$) and end to end distance as reaction coordinates, a) exp-CTD-non-phos, b) exp-CTD-5P-40P, c) exp-CTD-5P-22P-40P, d) exp-CTD-5P-12P-18P-32P and e) exp-CTD-5P-12P-18P-25P-32P-40P. In addition, $Y_1$-$Y_5$ represent a few of the lowest energy conformations of different CTD sequences.

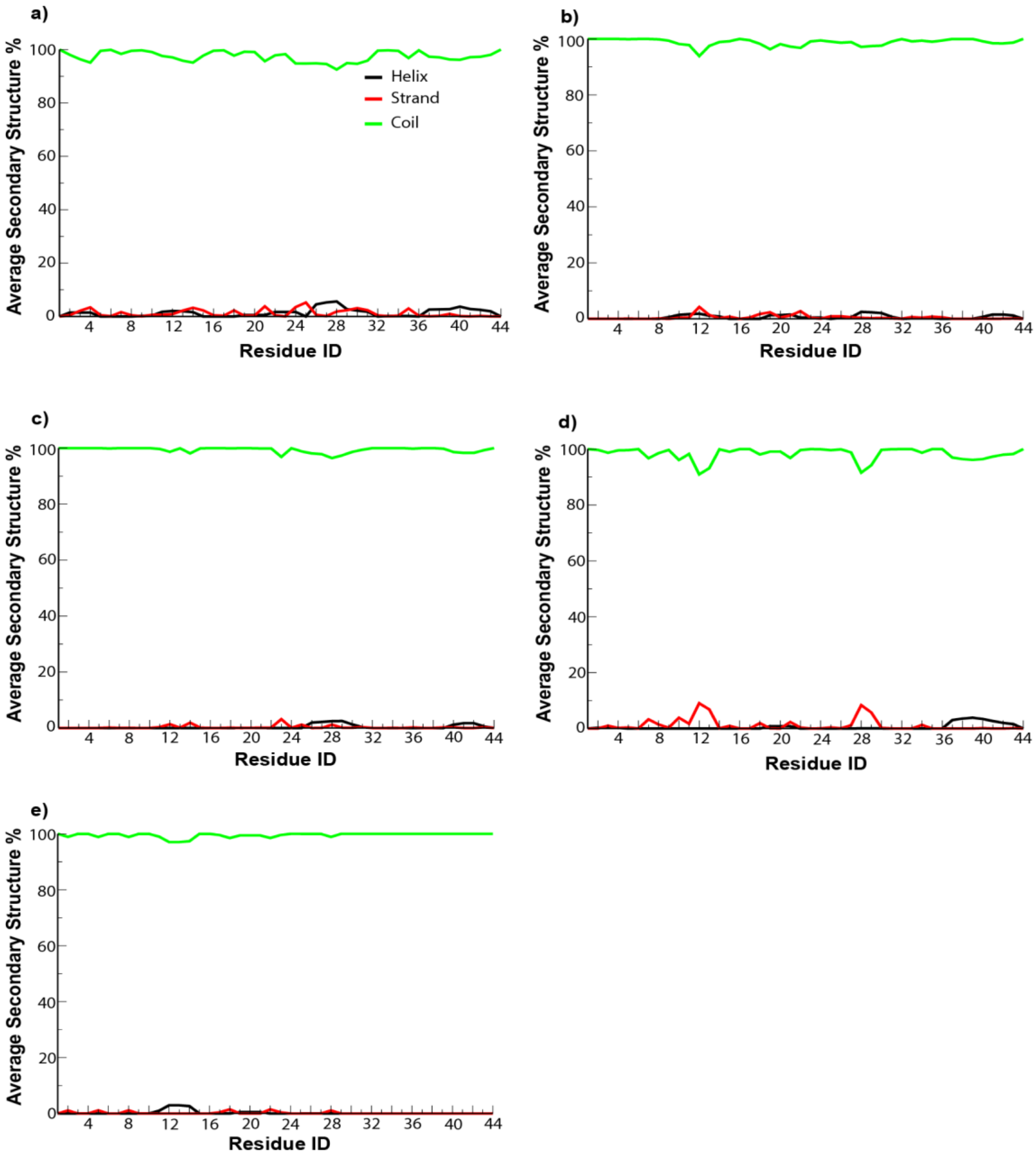


**Figure S5:** Average secondary structure percentages for CTD sequences with 44 residues. a) exp-CTD-non-phos, b) exp-CTD-5P-40P, c) exp-CTD-5P-22P-40P, d) exp-CTD-5P-12P-18P-32P and e) exp-CTD-5P-12P-18P-25P-32P-40P.

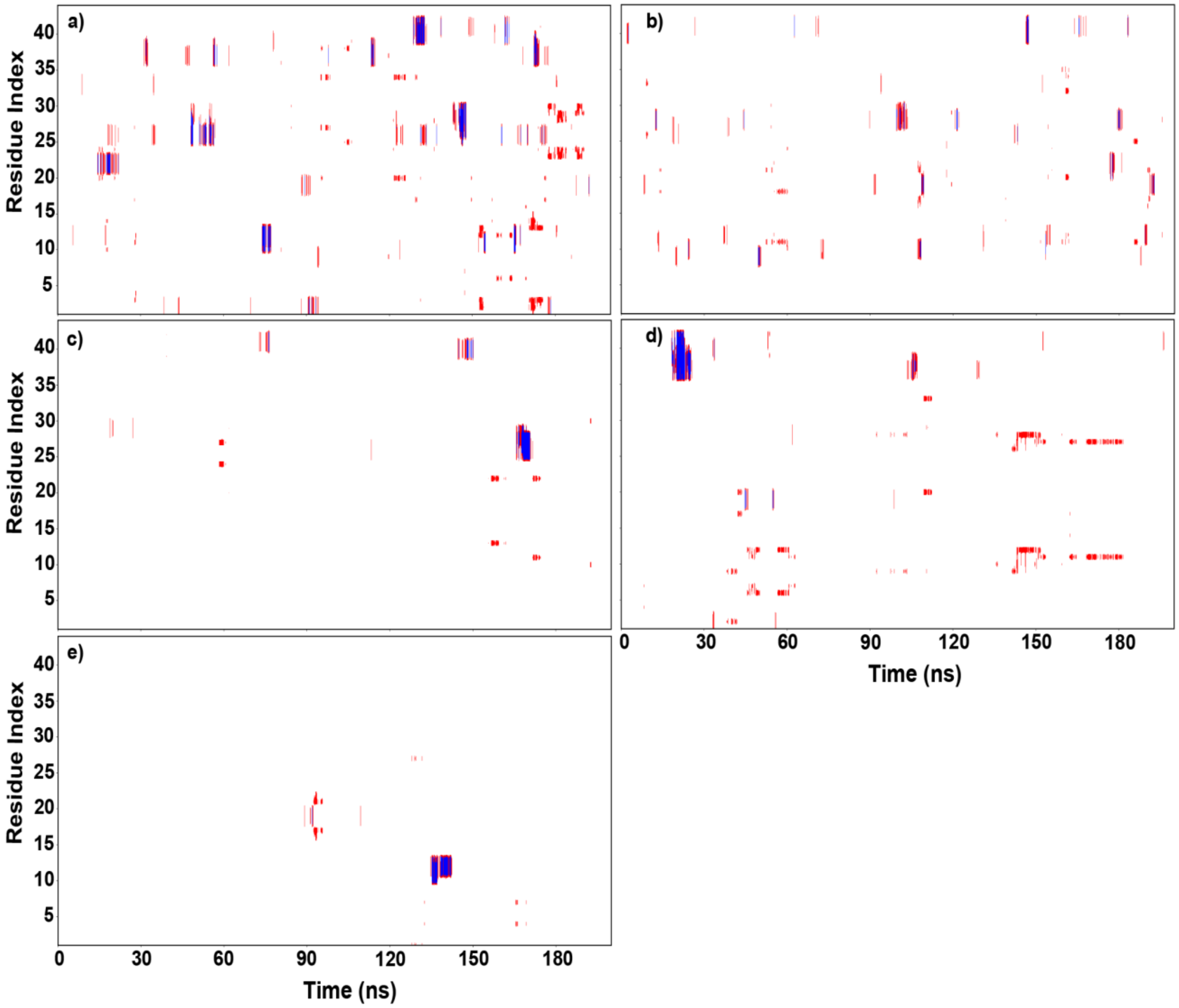


**Figure S6:** Secondary structure predictions with time for CTD sequences with 44 residues. Coil (loops, bends and turns) structures in white, helix (alpha helix, 3/10 helix and pi helix) structures in blue and strand (beta bridge and extended strand) in red. a) exp-CTD-non-phos, b) exp-CTD-5P-40P, c) exp-CTD-5P-22P-40P, d) exp-CTD-5P-12P-18P-32P and e) exp-CTD-5P-12P-18P-25P-32P-40P.

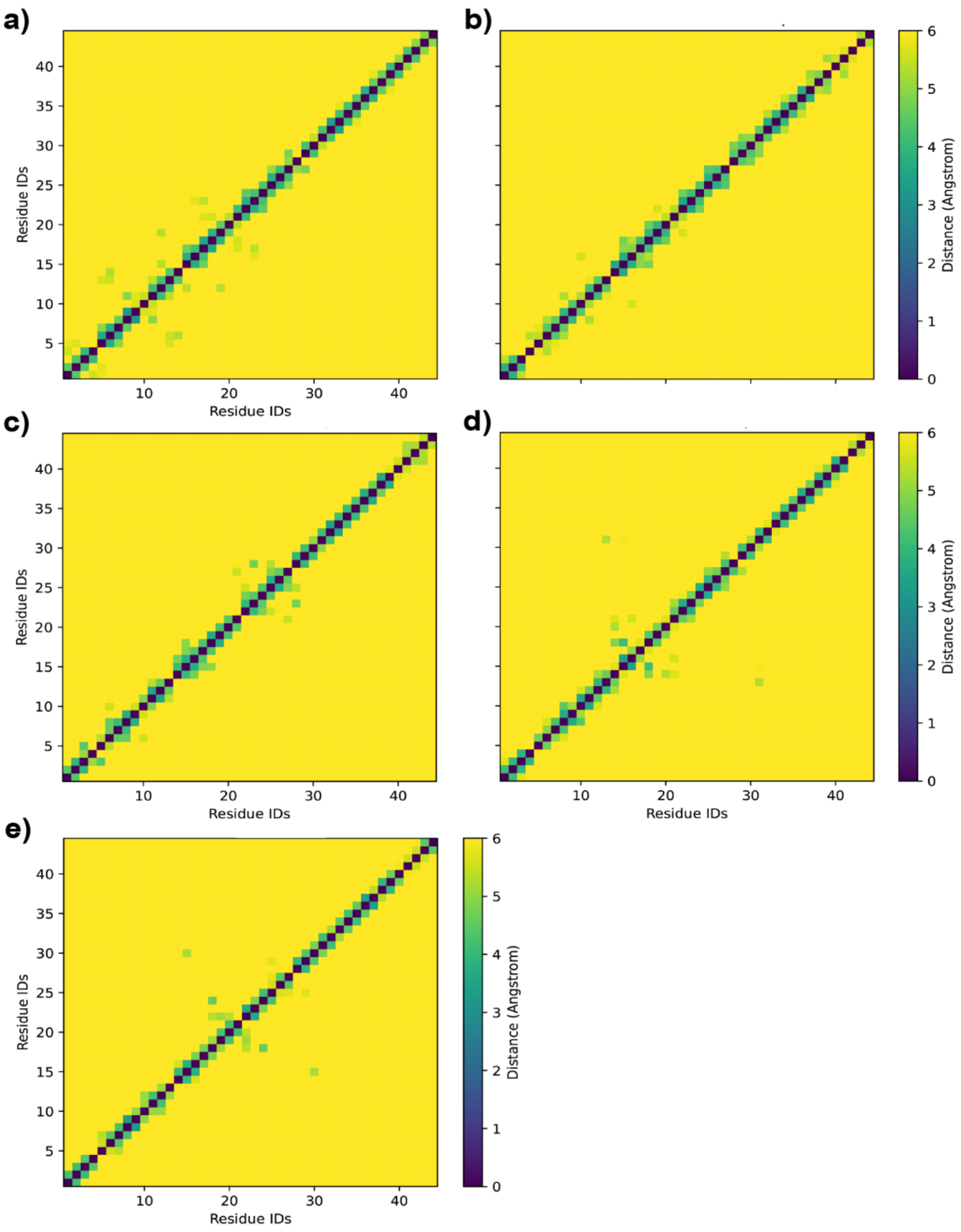


**Figure S7:** Distance maps generated from distances between center of masses of residues of CTD sequences with 44 residues. a) exp-CTD-non-phos, b) exp-CTD-5P-40P, c) exp-CTD-5P-22P-40P, d) exp-CTD-5P-12P-18P-32P and e) exp-CTD-5P-12P-18P-25P-32P-40P.

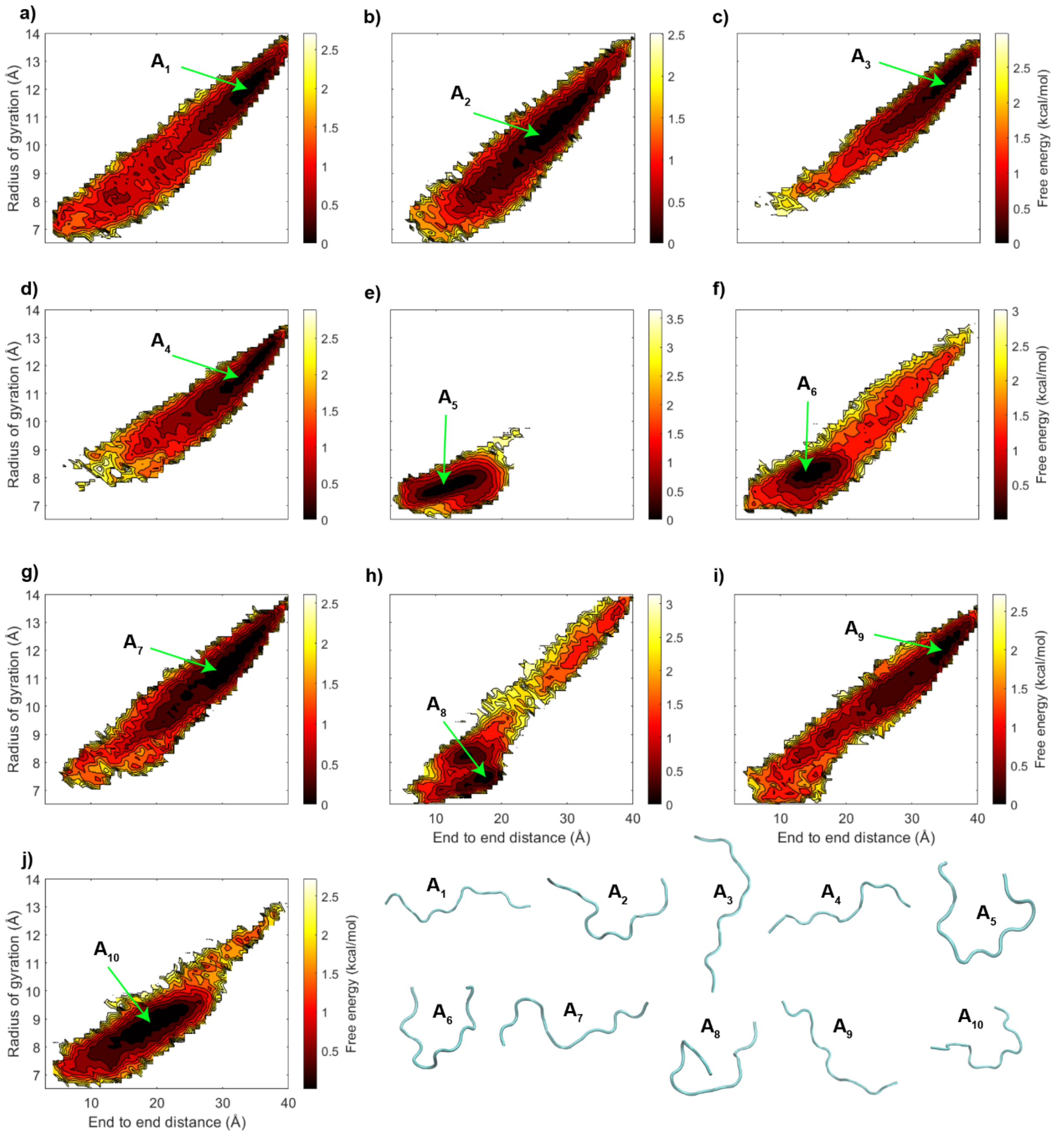


**Figure S8:** The free energy landscapes using radius of gyration ($R_g$) and end to end distance as reaction coordinates, a) 2CTD-non-phos, b) 2CTD-2P, c) 2CTD-2P-5P, d) 2CTD-2P-5P-9P, e) 2CTD-2P-5P-9P-12P, f) 2CTD-2P-5P-12P, g) 2CTD-2P-9P, h) 2CTD-2P-12P, i) 2CTD-5P and j) 2CTD-5P-12P. In addition, $A_1$-$A_{10}$ represents a few of the lowest energy conformations of different CTD sequences.

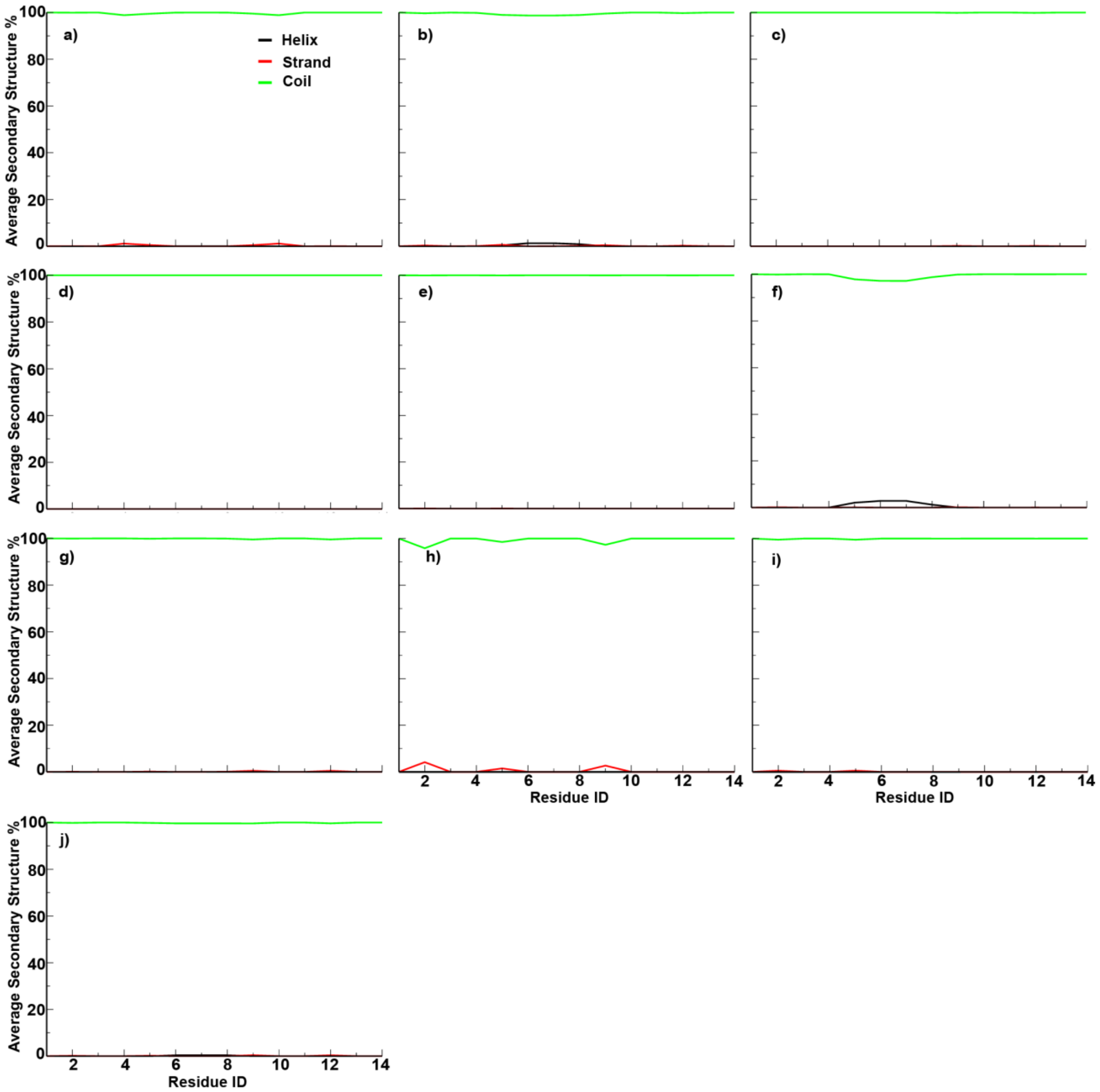


**Figure S9:** Average secondary structure percentages for CTD sequences with 14 residues. a) 2CTD-non-phos, b) 2CTD-2P, c) 2CTD-2P-5P, d) 2CTD-2P-5P-9P, e) 2CTD-2P-5P-9P-12P, f) 2CTD-2P-5P-12P, g) 2CTD-2P-9P, h) 2CTD-2P-12P, i) 2CTD-5P, j) 2CTD-5P-12P.

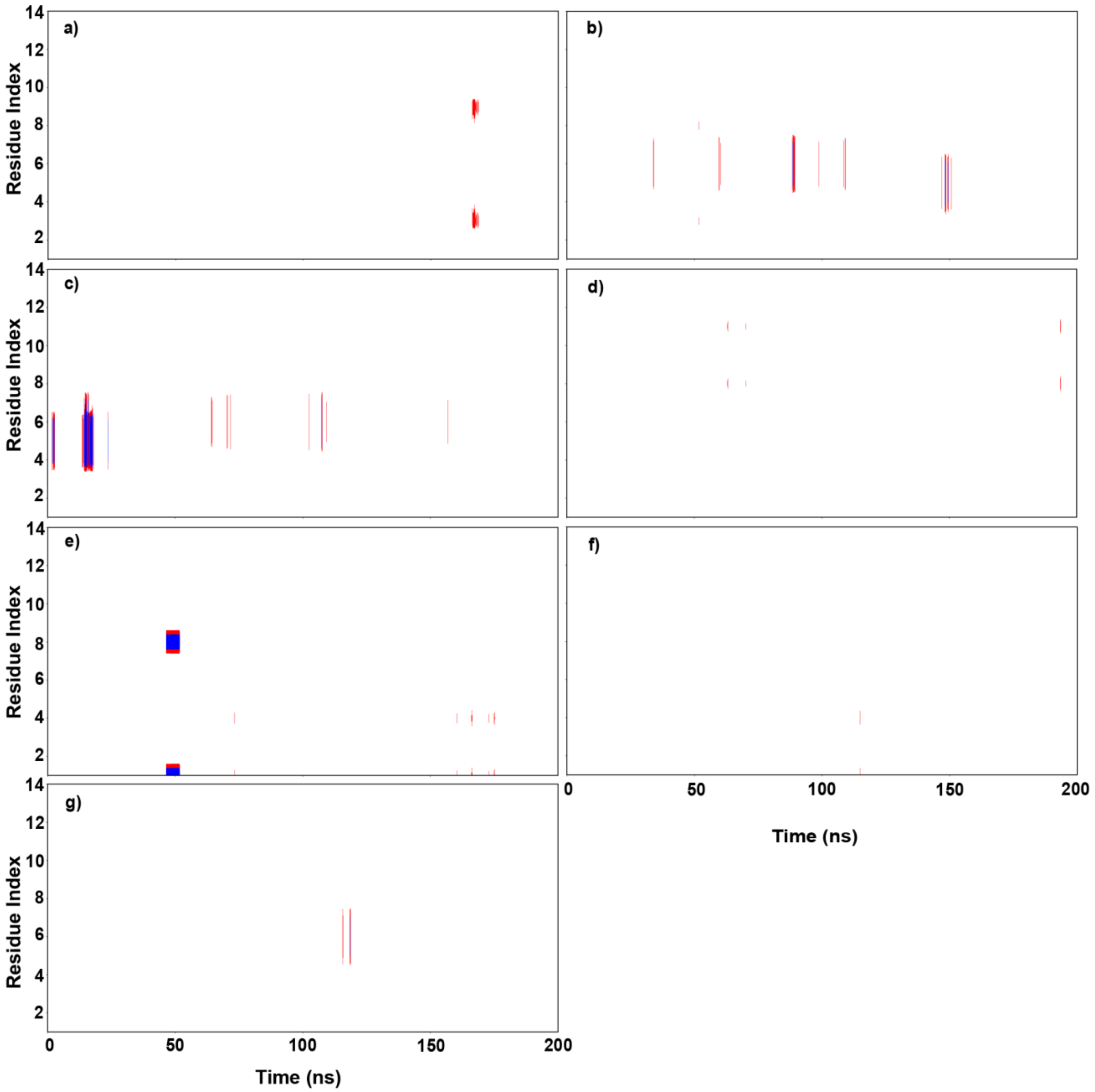


**Figure S10:** Secondary structure predictions with time for CTD sequences with 14 residues. Coil (loops, bends and turns) structures in white, helix (alpha helix, 3/10 helix and pi helix) structures in blue and strand (beta bridge and extended strand) in red. a) 2CTD-non-phos, b) 2CTD-2P, c) 2CTD-2P-5P-12P, d) 2CTD-2P-9P, e) 2CTD-2P-12P, f) 2CTD-5P and g) 2CTD-5P-12P. Moreover, 2CTD-2P-5P, 2CTD-2P-5P-9P and 2CTD-2P-5P-9P-12P are not shown in this figure because all the secondary structures were coils (loops, turns or bends) for those CTD sequences.

**Table S2:** Average number of intrapeptide H-bonds for CTDs with 44 residues and 14 residues.

| System | Average number of H-bonds |
|---|---|
| exp-CTD-non-phos | 2.8793 |
| exp-CTD-5P-40P | 2.9900 |
| exp-CTD-5P-22P-40P | 3.2640 |
| exp-CTD-5P-12P-18P-32P | 4.6228 |
| exp-CTD-5P-12P-18P-25P-32P-40P | 5.2206 |
| 2CTD-non-phos | 0.5363 |
| 2CTD-2P | 0.7627 |
| 2CTD-2P-5P | 0.8333 |
| 2CTD-2P-5P-9P | 1.5281 |
| 2CTD-2P-5P-9P-12P | 1.1204 |
| 2CTD-2P-5P-12P | 1.1054 |
| 2CTD-2P-9P | 1.2132 |
| 2CTD-2P-12P | 1.3214 |
| 2CTD-5P | 0.8317 |
| 2CTD-5P-12P | 1.1543 |

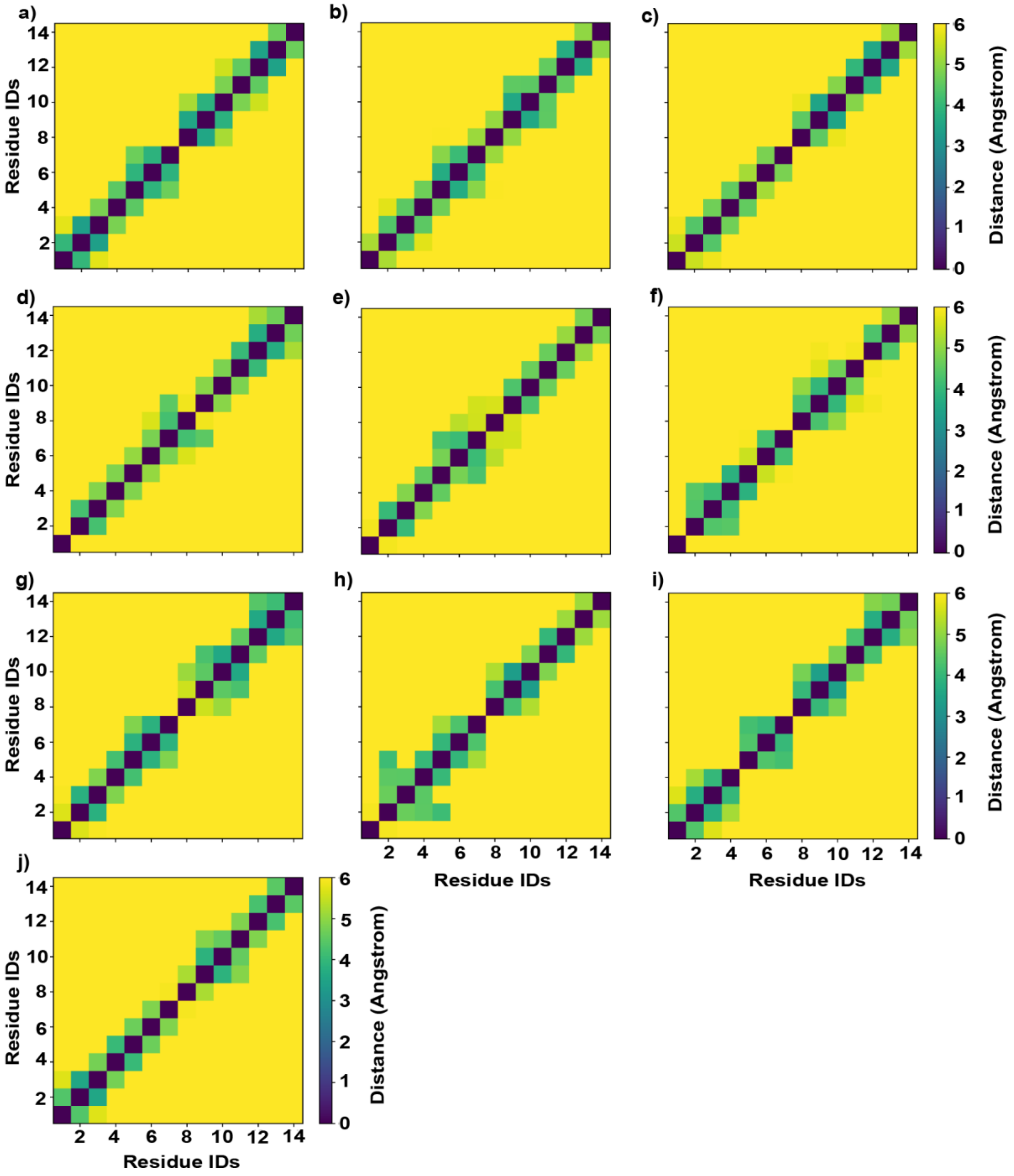


**Figure S11:** Distance maps generated from distances between center of masses of residues of CTD sequences with 14 residues. a) 2CTD-non-phos, b) 2CTD-2P, c) 2CTD-2P-5P, d) 2CTD-2P-5P-9P, e) 2CTD-2P-5P-9P-12P, f) 2CTD-2P-5P-12P, g) 2CTD-2P-9P, h) 2CTD-2P-12P, i) 2CTD-5P and j) 2CTD-5P-12P.

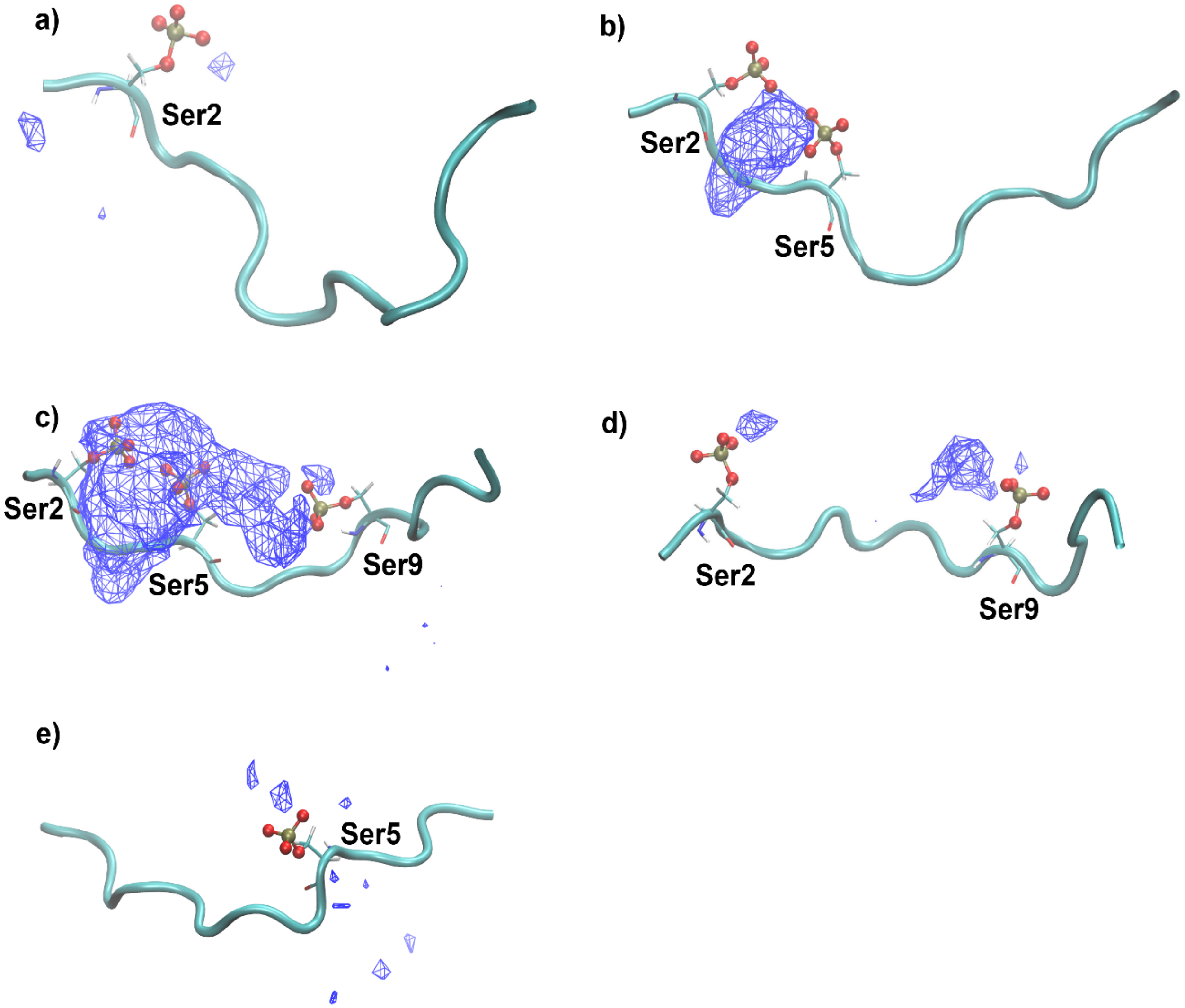


**Figure S12:** The average $Na^+$ ion density around the phosphate groups in serine (Ser) residues for the central structures of 2CTDs that did not contract. a) 2CTD-2P, b) 2CTD-2P-5P, c) 2CTD-2P-5P-9P, d) 2CTD-2P-9P and e) 2CTD-5P. The color codes are the same as Figure 6 in the main manuscript.
